# Mechanism-aware inference of response to targeted cancer therapies

**DOI:** 10.1101/2025.09.04.674143

**Authors:** Nilabja Bhattacharjee, Sreeram Chandra Murthy Peela, Abhishek Halder, Bernadette Mathew, Swarnava Samanta, Sakshi Gujral, Stuti Kumari, Smruti Panda, Ritwik Ganguly, Gaurav Ahuja, Subhajyoti De, Angshul Majumdar, Debarka Sengupta

**Affiliations:** Dept. of Computational Biology, Indraprastha Institute of Information Technology - Delhi (IIITD), New Delhi, India - 110020; Dept. of Computer Sciences and Engineering, Indraprastha Institute of Information Technology - Delhi (IIITD), New Delhi, India - 110020; Rutgers Cancer Institute of New Jersey, Rutgers, the State University of New Jersey, New Brunswick, NJ, USA; Dept. of Electronics and Communications Engineering, Indraprastha Institute of Information Technology - Delhi (IIITD), New Delhi, India - 110020; Infosys Centre for AI, Indraprastha Institute of0 Information Technology - Delhi (IIIT-Delhi), New Delhi, India - 110020

## Abstract

Targeted therapies like small-molecule inhibitors often work by blocking proteins that cancer cells rely on for survival. Omics based modeling of drug sensitivity alone lack mechanistic grounding. We propose **FORGE** (**F**actorization **O**f **R**esponse and **G**ene **E**ssentiality) a simple yet powerful joint matrix factorization framework that co-models drug response and target gene essentiality, enabling the stratification of promising treatment groups for targeted therapy consideration. **FORGE** also provides **Benefit Score** — a predictive score that estimates treatment efficacy from basal gene expression profiles. We validated the predictive performance of **FORGE** across multiple targeted therapies, including Erlotinib (EGFR inhibitor) and Daporinad (NAMPT inhibitor). Our meta-analysis of large scale *in-vitro* studies underscores **FORGE**’s ability to identify common determinants of drug vulnerabilities and target gene essentiality. Such convergences were not observed when treatment vulnerabilities and gene essentialities were modeled independently. We also demonstrated the universality of Erlotinib Benefit Scores by transferring transformations learned from high-throughput drug response studies across other published datasets, including the TAHOE-100M single-cell perturbation atlas and patient-derived xenograft studies. **FORGE** successfully identified key regulators within the molecular pathways targeted by these therapies, reinforcing its potential for mechanistically grounded treatment stratification.

## Introduction

Cancer is the second leading cause of mortality globally and is a complex disease affecting multiple organ systems [1]. Anti-cancer therapies typically target genes and pathways deemed essential for cancer cell survival, proliferation, or metastasis. However, gene essentiality varies significantly among patients and cancer types, limiting the efficacy of single-agent or combinatorial therapies [2, 3]. This lack of efficacy is further intensified by dose-limiting toxicities arising from on-target effects in normal tissues (e.g., EGFR inhibitors inducing dermatologic and gastrointestinal toxicity), which narrow therapeutic windows and negatively impact long-term outcomes [4, 5] Personalized medicine aims to circumvent these challenges by designing therapies using patient-specific omics profiles. This paradigm shift in cancer therapeutics has been facilitated by the availability of large-scale omics and high -throughput drug screens. Numerous databases like the Cancer Dependency Map (DepMap) [6], CREAMMIST [7], the Cancer Cell Line Encyclopedia (CCLE) [8], The Cancer Genome Atlas (TCGA) [9], and the NCI’s Patient Derived Models Repository (PDMR) (https://pdmdb.cancer.gov/web/apex/f?p=101:41) provide comprehensive genomic and functional profiling data for multiple cancer types. These resources were successfully employed by researchers globally in studying cancer mechanisms and identifying novel biomarkers for diagnosis and/or therapy.

Predictive machine learning (ML) or deep learning (DL) models utilize large-scale omics data as input features for predicting drug responses or identifying potential biomarkers for drug susceptibility. Numerous deep learning based models like CDRscan [10], DeepTTA [11], and DeepDR [12] utilize neural network architectures and aim to predict cancer drug responses using a variety of molecular and/or drug-related data. The CDRscan models drug properties encoded in its SMILES representation and mutational profiles of human cancer cell lines in a two-step convolutional architecture, while DeepTTA utilizes gene expression data along with drug SMILES information to predict cancer drug responses. DeepDR on the other hand, utilizes multiple omics layers - expression profiles, mutations (both SNPs and copy number variants) along with pathway enrichment scores (PES), and incorporate drug features as molecular fingerprints, SMILES and molecular graphs. All these models have shown promising accuracies in predicting drug responses. We previously developed and validated Precily, a deep neural network for inferring treatment responses using drug descriptors and gene expression profiles along with the PES [13]. PES aid in representing cells or samples as summarized values combining pathway activities and gene set enrichment scores [14]. In Precily, we discussed the issue of train–test data leakage, wherein reported accuracies of many deep learning models are artificially inflated due to the lack of proper cell line and drug blinding. Randomly splitting drug–cell line pairs across training and testing leads to the reuse of cell line expression profiles, enabling models to “memorize” baseline sensitivities and mimic the expected IC_50_ values for unseen drugs. To mitigate this, Precily enforced cell-line disjoint splits, ensuring that no cell line appeared in more than one dataset partition, thereby providing a more rigorous and clinically realistic evaluation of model performance.

Similar to drug response prediction, numerous models have focused on predicting gene essentiality used in drug discovery pipelines for various cancers. The DepMap database, in this context, forms the single large resource for providing gene essentiality data derived through functional assays using CRISPR-Cas9 knockouts [6]. Models like DeepHE [15] and XGEP [16] utilize gene expression data to infer gene essentiality among cancers. While XGEP can identify potential lncRNAs along with essential genes, DeepHE utilizes protein interaction networks (PINs) and various sequence features (gene length, GC content, etc.) in a deep learning framework. These models achieved an accuracy of 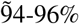 in predicting essential genes, and highlight the role of multiple genetic features in predicting gene essentiality. While the above discussed ML/DL models are gaining popularity, the practical applications of many ML/DL models are hindered by complexities in data integration, training, and model transferability to newer datasets [17]. These nonlinear approaches are particularly prone to overfitting, especially when trained on limited biological datasets. Deep neural networks, with their high parameter count, tend to “memorize” training data rather than learn generalizable patterns, resulting in poor performance on unseen samples [18–20]. This issue undermines model transferability and reduces clinical utility. Evidence suggests that pan-cancer, pan-drug ML models may not provide meaningful advantages over naive baselines, as a mean-baseline model that simply predicts the average IC_50_. This often yields predictive performance comparable to complex ML models, particularly in the challenging context of z-scored IC_50_ predictions where pan-drug ML models largely fail [21].

On the other hand, the dimensionality problem in traditional machine learning (*N* ≪ *P*) plagues numerous omics databases as the number of input features (genes, *P*) is substantially larger than the number of samples (patients or cell lines, *N*). Matrix factorization (MF) techniques, on the other hand, are resilient in handling such high-dimensional omics datasets, and play critical role in omics integration by linking latent embeddings to hidden biological states [22]. Techniques like Similarity-regularized MF (SRMF) [23], weighted graph regularized matrix factorization (WGRMF) [24] and Drug Sensitivity Prediction using Logistic Matrix Factorization (DSPLMF) [25] identify groups of cell lines that are susceptible to specific drugs. These methods learn *embeddings* from drug chemistry and cell line similarities, and can predict accurately the drug response values. Compared with deep learning and transformer-based models (as in DSPLMF evaluation) or 10-fold cross validation (in SMRF and WGRMF), these models demonstrated superior accuracy [23–25]. Matrix factorization models therefore provide a robust and interpretable alternative to the *black-box* nature of deep learning approaches.

In this study, we developed a matrix factorization–based algorithm called **FORGE** as an alternative approach to existing DL methods. **FORGE** uniquely relies on gene expression profiles—consistently shown to be the most predictive omics layer for drug response [26]—thereby reducing complexity while retaining predictive power. We leveraged publicly available large-scale resources, including DepMap (gene dependency and transcriptomics) and CREAMMIST (drug response profiles), to design **FORGE** as a joint-matrix factorization method to learn sample representations as a combination of latent weight matrices. Using the embeddings, we computed gene influence scores as contributions of each gene in predicting either dependency or IC_50_, and identified high correlation of these influence scores when using joint-modeling framework. We successfully showed that the learned embeddings are transferable, enabling accurate prediction of drug responses across both single-cell datasets and patient-derived xenografts. We further extended **FORGE** to derive a biomarker, the **Benefit Score**, which integrates the combined effect of drug response and target gene essentiality, thereby stratifying samples according to their likelihood of exhibiting favorable responses to targeted therapies. Furthermore, **FORGE** can detect key genes that impact therapy outcomes, positioning **FORGE** as one of the few computational models capable of not only predicting drug response but also generating biologically grounded hypotheses with translational potential for personalized cancer therapy.

## Results

### Overview of FORGE

Targeted therapy works by inhibiting key molecular vulnerabilities involved in cancer cell growth and survival. Large-scale pharmacogenomic studies such as GDSC [27], CCLE [8], and CTRPv2 [28] have generated vast amounts of high-throughput drug response data, while the DepMap project has produced systematic CRISPR-Cas9 dependency maps for many of the same cell lines. This overlap of datasets presents an unique opportunity to model drug response together with genetic dependency, so that drug predictions are grounded in mechanistic evidence of gene essentiality. By connecting pharmacological IC_50_ data with CRISPR-derived dependencies, we aim to align therapeutic response with underlying biology in a more robust and interpretable manner.

Matrix factorization–based frameworks, which decompose large data matrices into low-dimensional representations of features and samples, are well established in computational biology because they are simple, interpretable, and computationally efficient. We asked whether it is possible to jointly model target gene dependency and drug IC_50_ within such a framework, capturing shared biological structure while also preserving task-specific signals. To this end, we designed a joint matrix factorization model in which gene expression is mapped into a shared latent space through common weights, while task-specific transformations separately reconstruct dependency and IC_50_, optimizing general representation learning and task-specific prediction simultaneously.

In simple terms, **FORGE** begins with a gene expression matrix *G* ∈ ℝ^*n×p*^, where each row represents a cell line (*n* cell lines) and each column represents a gene (*p* genes). Instead of working in the very high-dimensional space of all genes, the model projects *G* into a smaller set of latent factors through a weight matrix *W* ∈ ℝ^*p×k*^. This gives us *Z* = *GW* ∈ ℝ^*n×k*^, where *Z* is the latent representation, and *W* is the encoder that learns which combinations of genes matter most. From this shared latent space, the model branches into two linear “heads.” One head predicts gene dependency, with weights *h*_*D*_ ∈ ℝ^*k×*1^ that are solved in closed form, and produces predicted dependency values 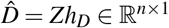. The other head does the same for drug sensitivity, with weights *h*_*I*_ ∈ ℝ^*k×*1^, producing predicted IC_50_ values *Î* = *Zh*_*I*_ ∈ ℝ^*n×*1^. The heads are updated by closed-form regression at each step, while *W* is updated by gradient descent. This alternation ensures that the two tasks remain linked through a common embedding while still refining the shared weights toward minimizing both prediction errors. In optimization terms, the gradient for *W* combines two parts: the error from the IC_50_ head and the error from the dependency head. A small 𝓁_1_ penalty is also added, encouraging sparsity in *W* so that only the most influential genes shape the latent factors. To make the outputs clinically interpretable, we defined a **Benefit Score** for each cell line. This is simply the difference between predicted dependency 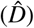 and predicted drug sensitivity *(Î)*. Intuitively, a high Benefit Score means the cell line is strongly dependent on the target gene (high 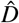) while also highly sensitive to the drug (low *Î*). Thus, this score acts as a per-sample measure of likely therapeutic benefit and allows us to stratify cell lines into those more or less likely to respond.

For drug sensitivity, we used the CREAMMIST database, which integrates *>*14 million dose–response measurements from five major pharmacogenomic resources under a Bayesian framework [7]. This harmonization overcomes inconsistencies between drug screening studies and provides robust, uncertainty-aware IC_50_ estimates for downstream modeling. For gene-level information, we relied on the Cancer Dependency Map (DepMap), which unifies omics profiles from the Cancer Cell Line Encyclopedia (CCLE) and genetic perturbation data from Project Achilles[29]. In particular, we used DepMap’s genome-wide CRISPR CERES dependency data, which systematically corrects for copy-number confounding and provides high-confidence gene essentiality scores, along with matched CCLE expression profiles for the same cell lines. Both resources span diverse tissue lineages, with CREAMMIST being dominated by pan-cancer, lung, and hematopoietic models (Supplementary Fig. 2a), while DepMap shows large representation of lung, lymphoid and skin cancers (Supplementary Fig. 2b). In total, 1,366 cell lines overlapped between DepMap and CREAMMIST, providing a harmonized, lineage-diverse cohort for joint modeling (Figure 1c). By combining DepMap’s standardized expression and dependency data with CREAMMIST’s harmonized drug response data, **FORGE** leverages the most comprehensive and mechanistically grounded resources available.

**Figure 1.**
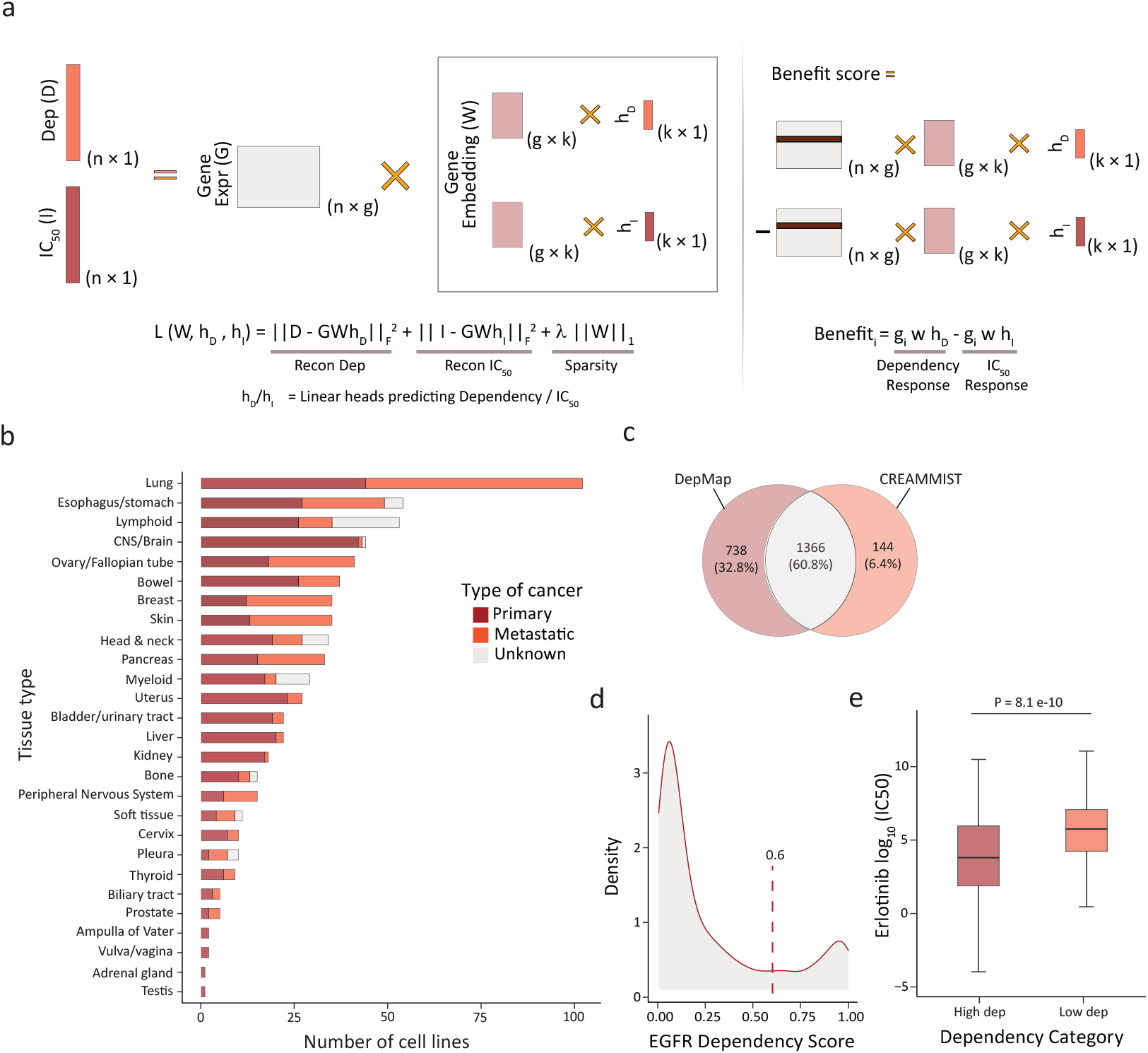
Latent representations of DepMap samples learned via joint-matrix factorization. **a**, Overview and implementation of **FORGE. b**, Tissue sources of the common cell lines (N=658) used for model building. **c**, Venn diagram depicting the number of cell lines unique and common to the DepMap and CREAMMIST databases. **d**, Distribution of EGFR dependency scores highlighting bimodality and the inferred cutoff for categorising samples. **e**, Boxplot showing the difference in erlotinib IC_50_ between samples with high and low EGFR dependency.

In the following sections, we present results from applying **FORGE** to specific drug–target pairs. Among such various pairs tested, EGFR dependency scores displayed a bimodal distribution, and erlotinib IC_50_ values segregated into two groups when stratified by median dependency (Figure 1d-e). Among these, 658 cell lines were common to both CREAMMIST and DepMap, with strong representation from lung, esophageal, and lymphoid cancers—providing a biologically relevant cohort to model the EGFR–erlotinib relationship (Figure 1b). We evaluate **FORGE**’s predictive performance, its ability to identify biologically coherent clusters of sensitive cell lines, and the interpretability of the gene influence maps that arise from the factorization. Together, these independent experiments demonstrate how a simple linear joint model can align genetic and pharmacological signals, offering both predictive accuracy and biological insight.

### FORGE successfully models EGFR targeted therapy

We sought to apply **FORGE** on large scale data to evaluate whether it could meaningfully stratify cancer cell lines according to their treatment sensitivity. While the framework can in principle be applied to any targeted therapy, we focused on Erlotinib, a small-molecule EGFR tyrosine kinase inhibitor that blocks downstream PI3K–AKT–mTOR signaling to reduce proliferation and increase apoptosis (Figure 2a). Erlotinib has been extensively studied and is clinically approved in non-small cell lung cancer (NSCLC) [30], as well as used in head and neck squamous cell carcinoma (HNSCC) [31], making it an ideal test case with both a well-characterized mechanism and clear clinical relevance. We hypothesized that a joint **FORGE** model would outperform independent models, since modeling dependency and IC_50_ separately risks learning two disconnected feature spaces, even though both are driven by overlapping cellular pathways. We also sought to confirm that the optimization procedure converges robustly across both tasks, given its alternation between closed-form updates and gradient descent, which could otherwise bias one prediction over the other. Finally, we asked whether the derived **Benefit Score** genuinely captures therapeutic potential by correlating with independent sensitivity stratifications, ensuring it represents a biologically meaningful metric rather than a modeling artifact.

**Figure 2.**
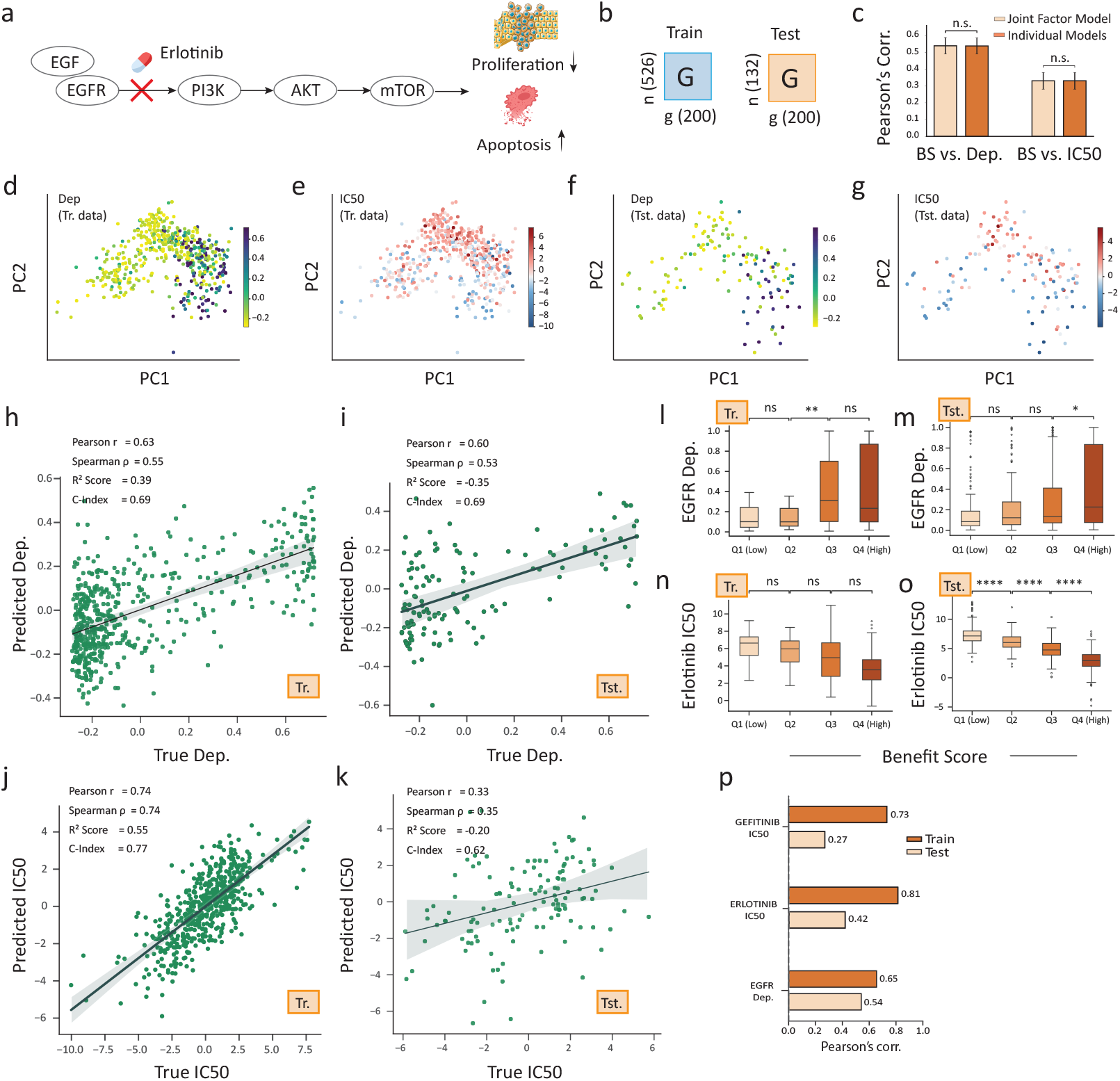
FORGE model performance evaluations. **a**, Mechanism of erlotinib activity in reducing cell proliferation and increasing rate of apoptosis. **b**, Dimensions of the input train and test datasets. The total number of samples are 658, and split in 70-30 ratio for model training and testing. **c**, Comparison of joint and individual matrix factorisation models. The Pearson *r* between Benefit Score and output features are shown. **d-g**, Projection of learned latent embeddings for training (**d-e**) and testing (**f-g**) datasets. A distinct clustering for high dependency and low IC_50_ are observed. **h-k**, Correlation between predicted and actual dependency and IC_50_ among the training and testing data. **l-o**, Categorising samples based on Benefit Score quartiles. **p**, Pearson *r* for multi-drug model for EGFR among the training and testing datasets.

To test these hypotheses, we trained **FORGE** on 658 cell lines overlapping between *DepMap* and *CREAMMIST*, using an 80/20 split (526 train, 132 test) (Figure 2b). Both dependency and IC_50_ predictions were modeled jointly in the shared latent space. In addition, we examined whether the *Benefit Score* aligned with independent decision variables, such as stratification across quartiles. Our results show clear spatial segregation between sensitive and resistant lines in the learned latent space: cell lines with high EGFR dependency and low erlotinib IC_50_ clustered distinctly, both in training and testing cohorts (Figure 2d-g). The model generalized well, with moderate to strong correlation between predicted and true values (e.g., Pearson’s *r* = 0.60 for dependency and *r* = 0.33 for IC_50_ in the test set), with the lower IC_50_ correlation likely reflecting the added complexity of drug action compared to the more direct effects captured by CRISPR-based dependency (Figure 2h-k). Sparsity in *W* acts as a built-in feature selector, pushing most gene weights to zero so that only the most informative genes remain. This yields a cleaner, more interpretable influence map, improves generalization by removing noise, and ensures the shared embedding focuses on genes contributing meaningfully to at least one task. We set the latent dimension to *k* = 40 based on prior evidence that the order of a few tens of components capture most variation in cancer expression data and represent key biological processes[32]. Empirical tuning showed that performance improved up to *k* ≈ 40 beyond which the train–test correlation gap increased, indicating overfitting(Supplementary Figure 2d). Importantly, the **Benefit Score** stratification aligned with biological expectations, showing higher values in more sensitive quartiles, confirming that it captured treatment-relevant information (Figure 2l-o). The joint model performed comparably to independent models in correlating the *Benefit Score* with outputs (Figure 2c); however, the correlation between their task-specific heads dropped significantly in individual models compared to the joint model, a point we will elaborate on in later sections.

Beyond the single-drug setting, we also explored a multi-drug extension with drug-specific heads, motivated by the idea that drugs targeting the same gene often act through shared mechanisms that should be captured within a common embedding. This design preserved a shared embedding to capture common biological mechanisms, while introducing drug-tailored readouts (*h*_*I*_ list) to account for differences in pharmacologic response across inhibitors. On the test set, this achieved correlations of *r* = 0.42 for erlotinib and *r* = 0.27 for gefitinib, both of which target EGFR (Figure 2p). Mechanistically, although both erlotinib and gefitinib are first-generation EGFR tyrosine kinase inhibitors (TKIs) that target the same receptor, their correlation differences showed significant differences. Exploring the datasets, we identified a few cell lines have uniform IC_50_ values for Erlotinib (Supplementary Figure 5a), and mean centering for these samples would produce values close to 0. We therefore presume that the high correlation in the EGFR-Erlotinib model could be due to this data artifact, rather than a true biological signal. Erlotinib is often administered at its maximum tolerated dose (MTD), leading to higher drug exposure in cell lines—especially those lacking canonical EGFR-activating mutations—which may enhance its predictive consistency [33]. Conversely, gefitinib, often given below its MTD and with a generally more favorable safety profile, may exhibit weaker signal-to-noise in some cell-line contexts, resulting in relatively lower predictive correlation [34]. In essence, the stronger performance for erlotinib versus gefitinib may reflects differential dosing potency and downstream signaling robustness, especially in heterogeneous biological contexts [35]. These results suggest that **FORGE** may be adapted to multiple drugs targeting the same gene, offering a scalable approach for broader applications.

### FORGE meaningfully clusters cancer cell-lines based on their treatment sensitivity

**FORGE** projects gene expression into a shared latent embedding, from which both dependency and IC_50_ are predicted. By integrating these outputs, the model derives a **Benefit Score** that reflects the potential therapeutic advantage of targeting a gene in a given sample. Correlating the latent space with the **Benefit Score** may provide a way to translate abstract model embeddings into clinically actionable sample groups. We asked whether samples sharing biological features would cluster together in the **FORGE** latent space, with such clusters revealing groups characterized by high EGFR dependency, low erlotinib IC_50_, and elevated **Benefit Scores**. Together, this would provide a chain of evidence linking the model’s latent structure to drug response and its underlying biology.

Continuing with EGFR-erlotinib system, we embedded all cell lines into the latent space *Z* = *GW* and applied hierarchical clustering with dynamic tree cutting (*cutreeHybrid* with *deepSplit* enabled and a specified minimum cluster size), a flexible approach that adaptively identifies both large and fine-grained groups without predefining cluster numbers. We chose hierarchical clustering because it does not require pre-specifying the number of clusters and provides a full dendrogram of sample relationships. When combined with Ward’s linkage, it identified compact and biologically coherent groups. The use of *deepSplit* allows detection of smaller, biologically relevant clusters that might otherwise be merged into broader groups. Cluster distributions were visualized using strip plots to capture separation of high- and low-sensitivity samples, and “sensitive clusters” were defined as those combining high EGFR dependency, low erlotinib IC_50_, and high **Benefit Scores**. We then performed differential expression between sensitive and non-sensitive groups, followed by pathway enrichment (*Enrichr-KG*), thereby linking model-derived clusters to gene-level programs and mechanistic pathways.

Dynamic clustering of cancer cell lines revealed multiple subgroups across the latent space, of which clusters 5, 6, and 19 emerged as the most “sensitive,” (Figure 3a,b). Swarm plots confirmed their distinct separation from the broader distribution of cell lines (Figure 3c,d). Among the 70 cell lines in these clusters, the majority were derived from head and neck squamous cell carcinoma (HNSCC) and non-small cell lung cancer (NSCLC). This is consistent with extensive evidence that EGFR is a key oncogenic driver in both HNSCC and NSCLC, where tumors are highly dependent on EGFR signaling for proliferation and survival.**FORGE** successfully recapitulates this biology by identifying these cancer types as the most sensitive groups, thereby validating the model’s ability to uncover clinically relevant dependencies and drug response patterns. Differential expression between these sensitive clusters and others yielded robust gene signatures, with upregulated genes including KRT17, KRT14, SERPINB3, ANXA6, and CXCR4—genes linked to epithelial integrity and keratinization. KRT17 has been shown to modulate EGFR expression through an integrin/Src/ERK/*β* -catenin–dependent pathway (e.g., in oral cancer models) [36], while SERPINB3 is implicated in epithelial integrity and EMT, affecting E-cadherin and *β* -catenin dynamics [37]. Notably, ANXA6, which was downregulated in the sensitive clusters, is a well-established negative regulator of EGFR signaling through receptor trafficking [38], reinforcing that **FORGE** captures biologically coherent mechanisms underlying drug response (Figure 3e). Enrichment analysis revealed strong associations with epidermis development [39], keratinocyte differentiation [40], keratinization, and skin development [41] (Figure 3f), processes that are well-established downstream of EGFR signaling, thereby reinforcing that **FORGE** captures biologically coherent pathways.

**Figure 3.**
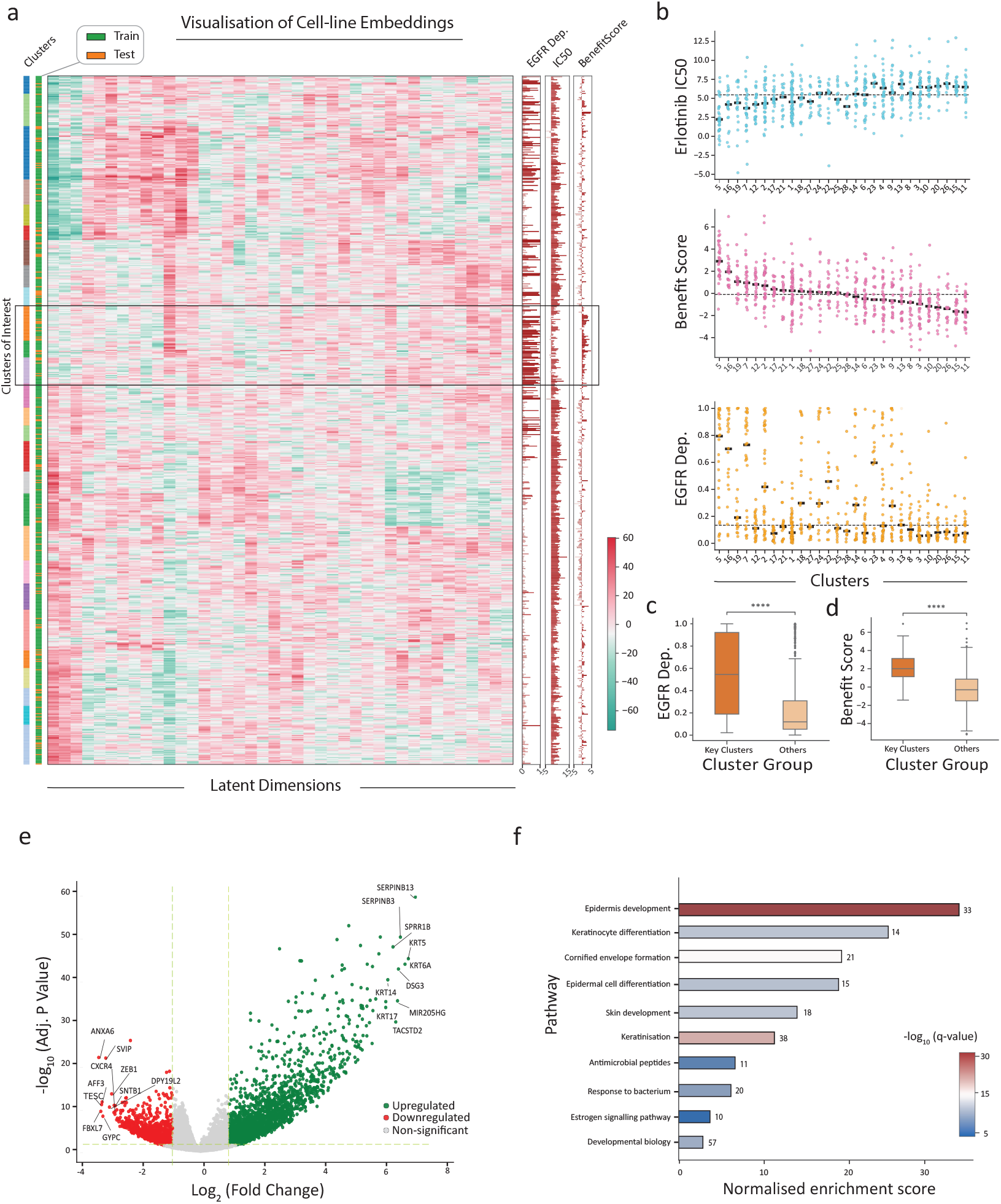
Analysing sample embeddings from the learned weights. **a**, Heatmap depicting the enrichment of latent dimensions in specific clusters. The left side color strips depict various clusters and distribution in the train-test categories, while the right side color strips depict EGFR’s dependency, erlotinib IC_50_ and inferred Benefit Scores for the samples. **b**, Swarm plot showing the top three clusters (clusters 5,6 and 19) with high Benefit Scores, low erlotinib IC_50_ and high EGFR dependency. **c-d**, Boxplots depicting the relative differences in EGFR dependency and Benefit Scores among the top three clusters and all other samples. **d**, Differential expression analysis (with top 10 significantly up and down regulated genes marked) of the key clusters as compared with other clusters. **f**, The enrichr-KG based GSEA analysis revealing key terms. The height of the bars represent the normalised enrichment score (Z-score) and are colored according to FDR q-values.

**FORGE** leverages latent embeddings and the **Benefit Score** to identify clusters of cell lines with high dependency and drug sensitivity, linking these groups to EGFR-related pathways. This demonstrates that the Benefit Score reflects biologically coherent signals, making it a practical and interpretable tool for stratification of samples by likely therapeutic response, with potential extension to patient stratification.

### Joint modelling of essentiality and response set common ground for model explanation

**FORGE** not only predicts cell-line–level outcomes but also decomposes the contribution of individual genes, providing task-specific effect sizes for both genetic dependency and drug sensitivity. For dependency, the **DependencyEffect** is defined as *Wh*_*D*_, where positive values indicate genes that increase model-predicted vulnerability to CRISPR knockout. For drug response, however, lower IC_50_ corresponds to greater sensitivity, whereas *Wh*_*I*_ naturally associates positive values with higher IC_50_ or, drug resistance. The model defines a **DependencyEffect** (*Wh*_*D*_) and an **IC**_50_**Effect** (−*Wh*_*I*_) for each gene, reflecting how strongly it influences vulnerability to CRISPR knockout versus pharmacological inhibition. Here, *Wh* takes a task-specific head (*h*_*D*_ or *h*_*I*_) and maps it back into gene space using the encoder *W*. This gives a vector of per-gene effect sizes, where each value reflects how much that gene contributes to the prediction. To align the interpretation of both tasks, we take the negative, defining the **IC**_50_**Effect** as −*Wh*_*I*_, such that positive values correspond to genes promoting sensitivity. This symmetry ensures that comparing **DependencyEffect** and **IC**_50_**Effect** highlights concordant biological drivers of both genetic dependency and pharmacological inhibition.

We hypothesized whether genes with positive effects on dependency (*Wh*_*D*_) would also negatively influence drug sensitivity (*Wh*_*I*_), reflecting shared biological drivers between CRISPR-based vulnerability and pharmacological inhibition. We further investigated whether discordant effects between these two axes could indicate mechanisms of drug resistance or pathway divergence. Finally, we tested whether the joint **FORGE** model improves this concordance compared to independent models, thereby offering a more interpretable view of gene-level influences across drug–target pairs.

To quantify gene-level contributions, we L2-normalized the task-specific heads (*h*_*D*_, *h*_*I*_) to obtain unit-norm vectors (*h*_*D*,norm_ = *h*_*D*_*/*∥*h*_*D*_∥, *h*_*I*,norm_ = *h*_*I*_*/*∥*h*_*I*_∥) of the EGFR-erlotinib model. L2-normalization removes scale differences between tasks while preserving vector direction, which encodes how latent features are weighted and is the relevant quantity for correlation-based concordance. Gene-level effects were then computed as **DependencyEffect** and **IC**50**Effect** computed using the unit-norm vectors. Plotting these values against one another allowed us to assess concordance between genetic dependency and drug sensitivity, with Pearson’s *r* quantifying global alignment. To highlight interpretable features, we extracted the top-20 and bottom-20 genes ranked by total influence, as these extremes are most likely to represent biologically relevant drivers or suppressors of EGFR signaling and drug response. Finally, we performed pathway enrichment analysis of these influential gene sets using *Enrichr KG*, to contextualize their functional roles and evaluate whether concordant effects reflect mechanisms. This pipeline was applied to both the joint and independent (dependency-only and IC_50_-only) **FORGE** models, and further repeated across multiple drug–target pairs to test generalizability.

Joint modeling produced a moderate alignment between these two axes (Pearson’s *r* = 0.20, *P* = 4.9 *×* 10^−3^; Figure 4a)), whereas individual models trained on dependency or IC_50_ alone showed much weaker concordance (Pearson’s *r* = 0.08, n.s.; Figure 4b)). This likely arises because the joint framework forces both tasks to rely on the same latent features, thereby reducing task-specific noise. We showed mathematically that this joint modeling approach can maximize the correlation between the dependency and IC_50_ effects for the genes under study (see supplementary note). Biologically, we validated our claim by testing across nine additional drug–target pairs where the joint model consistently yielded stronger gene-effect concordance than independent models (Repeated sampling t-test *P* = 6.28 *×* 10^−5^; (Figure 4c)). This supports the intuition that a shared embedding links dependency and pharmacologic response to common biological mechanisms.

**Figure 4.**
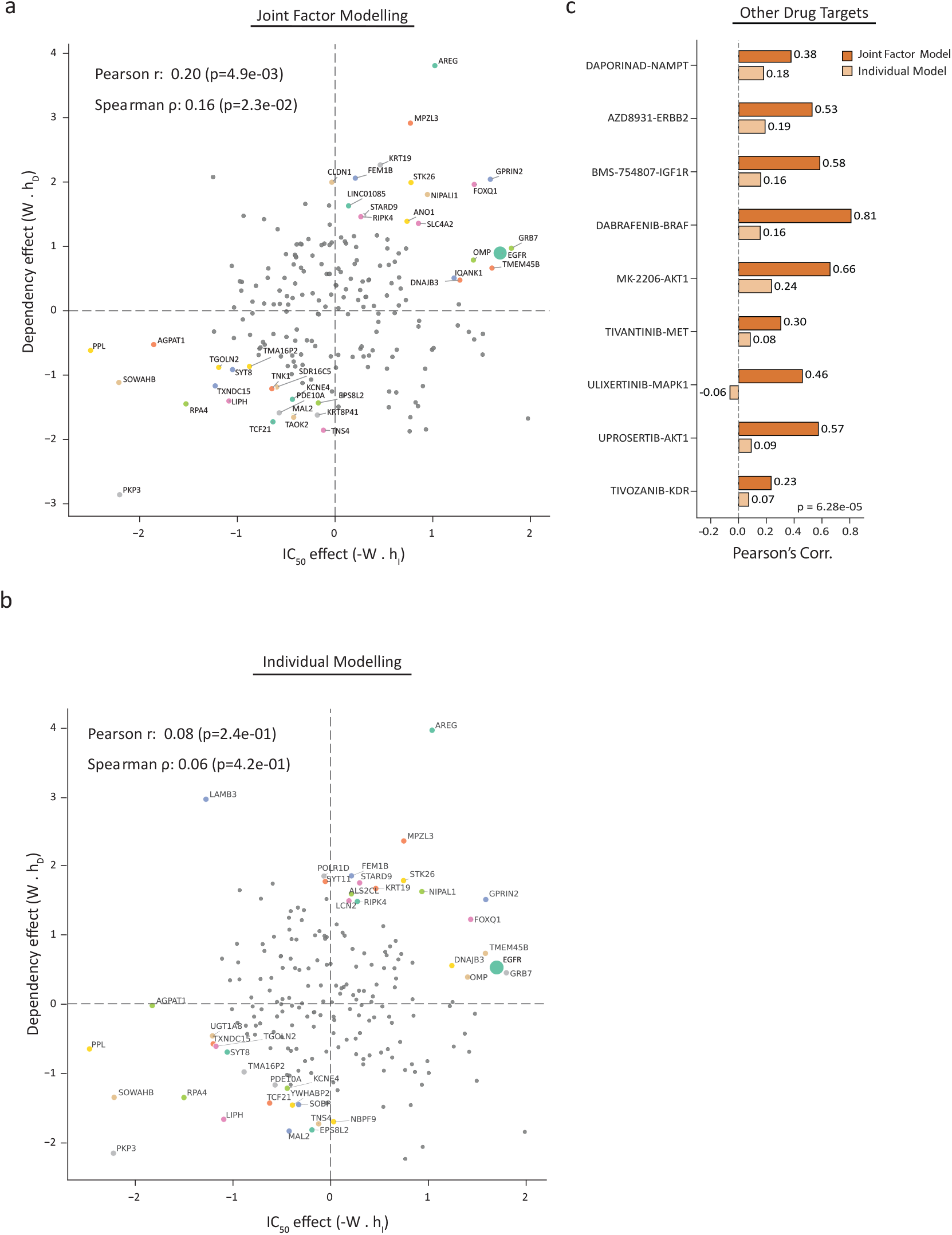
FORGE joint vs individual modelling. **a-b**, Scatterplot depicting the correlation between dependency and drug IC_50_ effects for each gene. **c**, Pearson *r* correlations between gene dependency and IC_50_ effects for various drug-target pairs with **FORGE** joint and individual models. Note: This figure illustrates sample-level latent embeddings derived from DepMap data using the **FORGE** model.

Biologically, the top 20 genes represent sensitizers—they promote both genetic dependency and drug response. These include AREG, GRB7, FOXQ1, ANO1, and MPZL3, all well-established enhancers of EGFR signaling or correlates of EGFR inhibitor sensitivity. For instance, AREG encodes an EGFR ligand whose secretion increases erlotinib response [42]; GRB7 [43] and FOXQ1 amplify EGFR pathway activity; ANO1 [44] stabilizes EGFR complexes at the membrane; and MPZL3 expression correlates with increased TKI sensitivity across cancers [45]. Together, these genes reinforce the idea that the model’s concordant quadrant highlights genuine EGFR biology. Conversely, the bottom 20 genes captured suppressors or resistance factors, including TNK1 and PPL. TNK1 disrupts Ras–MAPK activation downstream of EGFR, dampening its signaling [46]. PPL variants are linked to reduced erlotinib sensitivity in oral cancers [47]. Thus, the discordant genes represent biologically plausible modulators of resistance. Although 31 of the 40 top and bottom influencers overlapped between joint and individual models, indicating broad agreement on the most important genes, global concordance across all genes remained weak in the independent models. This suggests that while both approaches recover similar extremes, only the joint model aligns effects consistently across the genome, producing a coherent directional relationship between dependency and IC_50_. Enrichment of the top-20 and bottom-20 influential genes confirmed these patterns (Supplementary Figure 2e): terms included EGFR signaling and ERBB signaling, together with clathrin-mediated endocytosis [48], all directly tied to the EGFR receptor regulation and inhibitor sensitivity [49]. Notably, clathrin-mediated endocytosis highlights how receptor trafficking, beyond canonical signaling cascades, can shape erlotinib response by altering EGFR turnover and drug accessibility [50]. In addition, pathways such as keratinocyte differentiation and broader epithelial programs emerged, consistent with the tissue contexts (lung and head-and-neck cancers) where EGFR dependence is most pronounced [51]. These findings indicate that **FORGE** captures not only core EGFR signaling but also the regulatory and contextual mechanisms that influence drug response.

By aligning gene-level effects on both dependency and drug sensitivity, **FORGE** highlights coherent molecular drivers of response, including known EGFR pathway activators and resistance factors. This concordance strengthens biomarker discovery by ensuring that influential genes are not task-specific artifacts but instead reflect shared biological mechanisms.

### Benefit score generalizes to independent *in vivo* and *in vitro* studies

We demonstrated the accuracy of **FORGE** through a critical dissection of the model and its various components. Through a thorough investigation of the predicted IC_50_ and gene-level dependencies, we established that the shared latent embeddings coupled with simple baseline gene expression can reveal potential differences among samples, and described the key genes that drive the model predictions. Similarly, we computed Benefit Scores as a metric to detect ‘susceptible’ clusters. While the more popular deep learning models are supposed to generalize well, they often fail due to memorization and learning inherent ‘noise’, especially in smaller datasets [52]. On the other hand, **FORGE**, being a linear model, has fewer degrees of freedom relative to deep learning networks. Furthermore, the sparsity regularization included in computing the loss function (Equation 6) reduces the risk of overfitting.

We tested the generalization capabilities of **FORGE** in identifying probable **high** responders to a targeted therapy given baseline gene expression values. To this end, we applied the Benefit Scores estimated using the shared latent embeddings (Equation 13) and publicly available gene expression datasets - the Tahoe 100M and the PDX datasets - both of which form the external validation datasets in the present study. The validation datasets belong to two different biological models - the Tahoe-100M data represent cell-line perturbation effects at single-cell level, while the PDX dataset contains baseline measurements of tumors in mouse models. Through this analysis, our aim was to re-verify our initial hypothesis that samples with higher Benefit Scores have more favorable responses to therapy. This critical step of validating our model using external datasets highlights **FORGE**’s generalization capabilities and highlight possible effects of differences in distributions of training and testing data.

#### FORGE categorized cell-line specific response strengths in perturbed scRNAseq

The Tahoe-100M represents a large resource of single-cell expression profiles of 49 + 1 cancer and normal cell lines perturbed with erlotinib (at varying concentrations) and DMSO (as a control) (Figure 5a). The available expression (raw counts) of 62,710 transcripts and corresponding metadata were retrieved as AnnData objects for each concentration category, which were then aggregated into pseudo-bulk counts per cell-line, and normalized using Limma’s Voom method (Figure 5a). For this purpose, we considered DMSO-treated cells from plate 9 (**high-concentration**) as the control, and normalized the drug-treated pseudo-bulk counts. Converting single cell data into pseudo-bulk counts is a standard practice for differential expression analysis, and aims to overcome the inherent sparsity associated with single cell transcriptomic profiles.

We ensured to use baseline expression profiles from DMSO-treated cells to compute Benefit Scores (as stated above), and prevent any bias due to drug-induced perturbations leaking into the model predictions. Among the 200 HCGs used in model training, one (0.5%) gene (SEPT7P6) was missing in the Tahoe-100M dataset, and two genes (CTGF and TMEM8A) were encoded using their aliases (CCN2 and PGAP6 respectively). Recoding these genes into their aliases and using gene weights derived from the original latent embeddings from the DepMap dataset, the computed Benefit Scores had a median of 5.44 across cancer cell lines (Supplementary figure 6E). In sharp contrast, the normal cell line (hTERT-HPNE) exhibited a markedly negative score (–7.34), highlighting a clear separation between malignant and non-malignant contexts. This differential suggests that the oncogenic background in specific cancer cells confers a substantial gain in susceptibility. Following this observation, we wanted to identify the transcriptomic shift associated with therapy (at varying concentrations) among these score categories.

**Figure 5.**
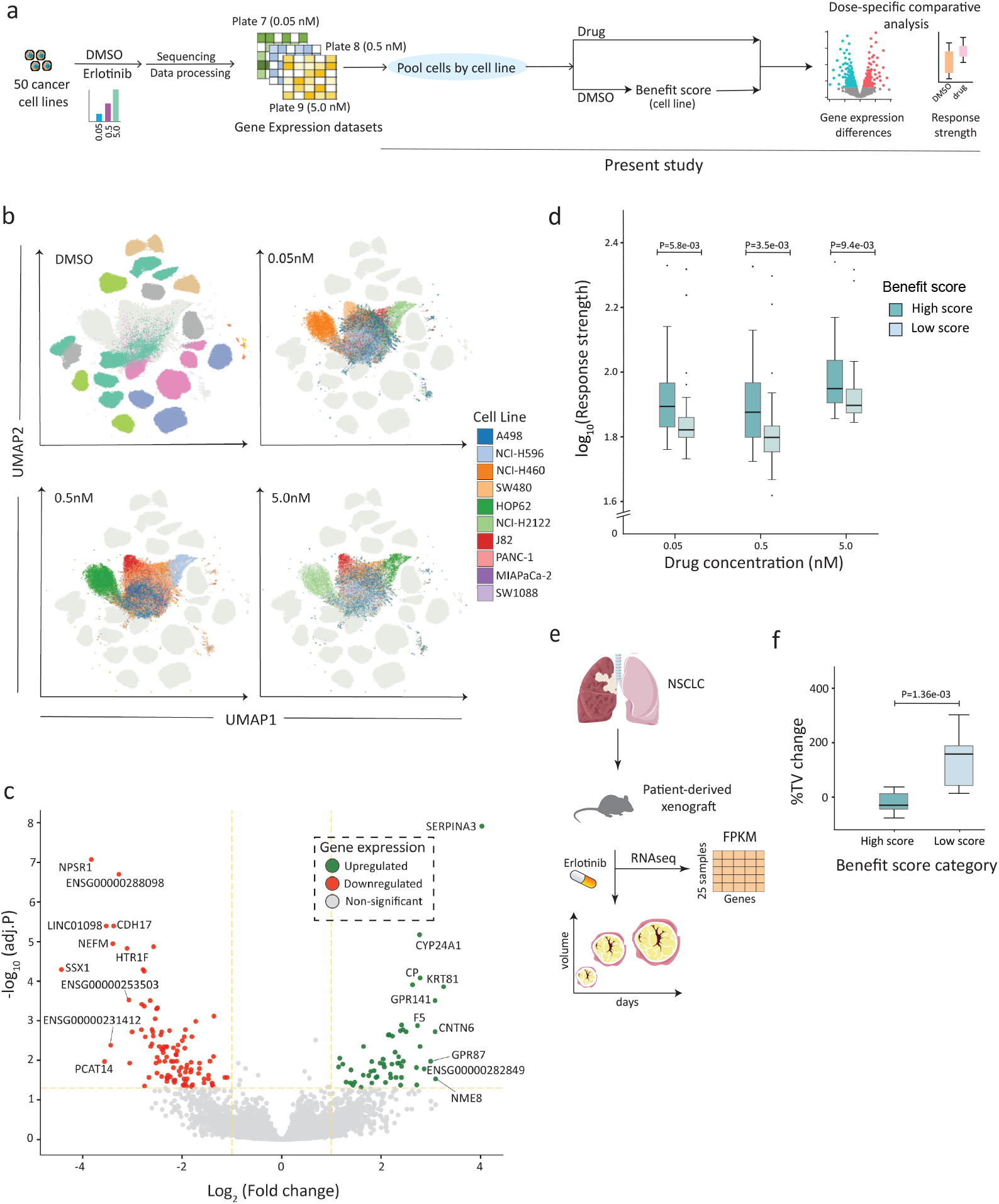
Validating **FORGE** model using Tahoe-100M (**a-d**) and PDX (**e-f**) datasets. **a**, Overveiw of Tahoe-100M experimental setup and pre-processing. **b**, UMAP plots depicting single cells for the best responding cell lines when treated with DMSO or erlotinib (at various concentrations). **c**, Differential gene expression analysis of all the cell lines based on Benefit Score categories. The top 10 significantly up/down-regulated genes are labeled. **d**, Box-plot depicting response strengths at varying concentrations among low and high scoring groups. **e**, Overview of experimental and data collection methods for PDX dataset as reported in the original publication. **e**, Box-plot showing the difference in percent change in tumor volume (%TV Change) between high and low Benefit Score categories at T=15 days. Note: All the P-values were computed using one-sided Mann-Whitney test.

We restricted our analysis to a common set of 17,667 genes between DMSO and drug-treated cell lines for quantifying the changes in transcriptome profiles. The response strength acts as a proxy for the shifts, and were computed as the Euclidean norm of the overall distance between the expression vectors belonging to treated and untreated samples. Across all the concentrations, ten cell lines showed predominant transcriptomic shifts (Figure 5b), and the response strengths were dose-dependent with greater shifts observed at higher concentrations (5.0 nM, plate 9) (Supplementary figure 6F). Interestingly, no marked differences were observed at lower drug concentrations (0.05 nM and 0.5 nM), indicating a possible drug concentration threshold that can elicit observable transcriptomic reprogramming. When stratified by Benefit Scores, we observed a dose-independent marked increase in response strengths among cell lines with higher Benefit Scores (Figure 5d). These findings demonstrate that the Benefit Scores capture a more underlying heterogeneity of responses that are not driven through drug concentrations, and highlight its utility as a potential biomarker for detecting samples that can have marked therapeutic responses.

Our observations indicate that the transcriptomic reprogramming is a dose-dependent feature, while Benefit Scores capture the underlying biological diversity that can enhance the shift. To further understand the biological mechanisms and pathways that are different among samples with higher benefit scores, we performed a differential gene expression analysis focusing on eight (16%) cell lines with maximal (in either directions) response strength. We used the baseline (DMSO-treated) expression values only to avoid any changes induced through drug challenge from percolating into our analysis. With DESeq2 and cell-line labels and concentration categories as additional co-variates, we identified a set of 158 (0.26%) differentially expressed genes (figure 5c). These genes were associated with various pancreatic cancer subtypes (WikiPathways WP5390) and gastric cancer. When we applied more permissive thresholds for enrichment analysis (minimum number of mapped genes = 3 and minimum number of terms per gene = 1), additional pathways related to metaplasia and epithelial maintenance emerged(Supplementary Figure 6G), affirming our use of Benefit Scores in capturing key pathways and genes that can influence drug susceptibility.

#### FORGE identified samples with beneficial tumor responses in mouse xenograft models

The PDX tumor samples from Gao *et al*. consisted of 399 PDX models, among which 25 were NSCLC samples treated with erlotinib. The dataset contains FPKM values estimated through read mapping with GRCh37 as the reference genome. As we employed voom normalization for all the earlier datasets, we converted FPKM values to pseudo-raw counts using a customized reverse-engineered formula with two hyper-parameters—the average number of mapped fragments (in millions) and the read length (equation 3). Given that the data were generated using paired-end mode with read lengths (*L*) ranging from 75–100 bp, we estimated the fragment size as 200 bp and the effective length as *L*− 199. We used an expected number of mapped fragments as 35 million, following our observation that the usual number of fragments available in NCBI’s SRA database ranges between 20–35 million (https://www.ncbi.nlm.nih.gov/sra). Though our above hyper-parameters are not exactly derived from the original source, using a range of read lengths from 75 bp to 100 bp and 20 to 50 million fragments, we observed a small shift in the log_10_(pseudo-raw counts)(see Supplementary Figure 7a). We subsequently used L_30_ model for validating this dataset.

Among the two response variables studied, we used the %TV Change, a measure of change in tumor volumes at different time-points when compared with original tumor size, as the key response metric. Identifying relative sparsity of these observations beyond 20 days for NSCLC tumors, we restricted our analysis to 15^th^ day by binning data-points between 10^th^ and 15^th^ day and summarized using mean. The pseudo-raw counts were normalized using Limma’s Voom normalization after removing lowly expressed (N=6817, 34.7%) genes and sub-setting with the original HCG list, leaving 130 genes for computing Benefit Scores. For computing Benefit Scores, we tested models with latent dimensions of 30 (L_30_ model) and 40 (L_40_ model) by considering the gene weights computed from the respective **W** and **h** matrices. We performed this analysis exclusively for this dataset as it contains actual tumor responses as %TV Change, and we believe this represents a real-world scenario for applying Benefit Scores. It must be noted that we identified substantial correlation (Pearson r = 0.74, *P* = 2e-35) but the R^2^ was 0.548 among the gene weights for both the models, revealing a similar trend but slightly higher weights from the L_30_ model (Supplementary Figure 7d).

At T=15 days, we identified a substantial (one-sided Mann-Whitney *P*=1.36e-3) reduction in tumor volumes among samples from high score group (median = -29.9, IQR = -44.43, 12.8) when compared to those from low score group (median = 158.26, IQR = 44.03, 190.09) (Figure 5f). It must be noted that in this variable, the negative values represent tumor repression as the volumes decreased when compared with the baseline (T=0 days). Across different time-points, we similarly observed a generalized patterns of tumor repression among high scoring samples (Supplementary Figure 7e). However the initial responses at T=5 days were not different among the two groups, indicating a slower effect of drug in tumor regression. Similarly, the BestAverageResponse (patient-wise averaged percent change in tumor volumes at *T* ≥ 10) was also lower among the samples with higher Benefit Scores (Supplementary Figure 7f), indicating a favorable outcomes within this cohort. This metric was followed in the original publication, and it summarizes the speed, strength and durability of response, and only the samples with higher Benefit Scores belong to mPR (partial response) while those among low scores were mPD (progressive disease) following mRECIST criteria (Table 1). Owing to the low sample size (N=25), we failed to observe any significant differences in proportion of favorable outcome (mPR) (Chi-squared P=0.1), but the effect was moderate to strong (Crammer’s V = 0.429). These observations together indicate our Benefit Scores were able to capture the tumor response trends by accurately identifying the cohort with favorable outcomes.

**Table 1.** Response categories (mRECIST) among high and low scoring PDX models.

| mRECIST category | High score | Low score |
| --- | --- | --- |
| mPD | 5 | 9 |
| mPR | 3 | 0 |
| mSD | 5 | 3 |
| Chi-squared statistic | 4.610 |  |
| p-value | 0.100 |  |

The above criteria, derived from [74] and based on BestAverageResponse, are as follows: mPD – Progressive Disease (tumor volume increased >30%); mPR – Partial Response (tumor volume reduced ≥20%); mSD – Stable Disease (tumor volume change <30%). None of the 25 samples showed complete response (mCR, >40% reduction). All definitions follow the original RECIST criteria as described in [74].

We identified 70 (35%) genes from the original HCG list which were missing in the PDX dataset. These genes had no potential aliases or changes in HGNC symbols, and therefore failed to map across genome builds (GRCh37 and GRCh38). This huge number of missing genes may have substantial influence on computing the Benefit Scores, and was specifically isolated to this dataset. We observed no significant difference of gene weights among the genes ‘missed’ in the PDX dataset (Supplementary Figure 7c). Similarly, 11 of the missing genes were also a part of the top 40 highly influential genes, indicating a potential impact on model predictions and the estimated Benefit Scores. In fact, these 11 genes were associated with epithelial-mesenchymal interactions in lung and kidney development (TCF21), cell growth and apoptosis (STK26), DNA repair (RPK4) or tumor suppression (LINC01085), to name a few. It has been observed that the over-expression of LINC01085 can be influential in treating hormone-independent prostate cancer that is resistant to docetaxel [53]. While we agree that much information has been lost due to these missing genes, we believe the sample-level encodings through Benefit Scores are still comparable.

Through a detailed analysis of our external validation sets, we proved the utility of Benefit Scores in understanding the transcriptomic re-programming (Tahoe-100M dataset) and in predicting favorable responses to therapy (PDX dataset). The gene signatures among Tahoe-100M data represented cancer-related pathways, making our scores biologically meaningful. We also demonstrated the role of Benefit Scores in identifying a highly sensitive cohorts - at cell-line level (DepMap data) and the PDX models, and validity of our **FORGE** model by transferring the weights across datasets. Given the substantial performance of our model, we now extend it to other drug-target pairs in the subsequent sections.

### FORGE predictions remain consistent across other drug-target pairs

Having validated **FORGE** on the well-characterized EGFR–erlotinib axis, we next turned to other drug–target pairs to build confidence in its broader applicability. By implementing the framework beyond a single setting, we aimed to explore whether the Benefit Score and latent embeddings consistently recover biologically expected treatment patterns across diverse pharmacological contexts.

We hypothesized that the Benefit Score would align with tissue lineage, such that model-derived scores recover tissue-specific drug sensitivities consistent with known biology and clinical evidence. This would demonstrate that the score not only predicts response but also reflects established treatment patterns. We further asked whether drug–target pairs achieve pharmacological separation in the latent space, as this would indicate that **FORGE** embeddings capture drug-specific biology rather than generic expression structure. In addition, we tested whether **FORGE** maintains predictive accuracy on both dependency and IC_50_ tasks across multiple pairs, to evaluate robustness of its multitask design. Finally, we examined whether **FORGE** can reproduce mechanistic insights for targets beyond EGFR–erlotinib, since this would show that the framework generalizes to uncover interpretable biology across diverse contexts.

We selected ten drug–target pairs with sufficient overlap in cell lines. For each pair, we computed the **Benefit Score** for every cell line as the difference between the predicted dependency 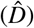 and the predicted IC_50_ (*Î*),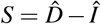. Lineage-specific patterns were visualized using heatmaps of **Benefit Score**, and latent embeddings were projected into UMAP to assess pharmacological separation. Each model was trained with 10 bootstrap replicates, using random seeds defined as (42 + *b*) for *b* = 0–9, to systematically evaluate stability. We presented our results for the NAMPT-Daporinad, and identified ‘susceptible’ clusters through methods described earlier. Differential expression between sensitive and other lines was performed using **limma**, which applies linear modeling with empirical Bayes variance shrinkage to increase power in high-dimensional settings. Enrichment of the resulting gene lists was carried out using *Enrichr-KG*. Finally, lineage–drug matches were benchmarked against published clinical data.

Comparing Benefit Scores derived from different models, we identified trends as expected for specific tumor types. The benefit scores had a clear lineage-specific patterns across drug–target pairs (Figure 6a), and many drug-target pairs shared a specific latent space (Figure 6b). The Dabrafenib–BRAF pair, a preferred treatment in melanoma, achieved the highest scores in skin lineage cell lines [54]. Similarly, AKT and MET inhibitors, commonly used in lymphoma, showed strong activity in lymphoid and myeloid lineages, consistent with clinical trials of these agents in lymphoma [55] and AML [56]. Similarly, the Daporinad–NAMPT pair produced high scores in hematological malignancies where it is effective [57]. Interestingly, we identified a novel link between Tivozanib (KDR), a highly selective VEGFR tyrosine kinase inhibitor, and myeloid cancers. Evidence suggests high expression levels of VEGF and its receptor VEGFR-2 among AML patients [58], and VEGFR-2 is selectively expressed on tumor-associated myeloid cells [59]. UMAP plots demonstrated partial pharmacological separation across pairs, though many models clustered in a central, mixed region, indicating both expected lineage segregation and shared biology across tissues (Figure 6b). Bootstrap runs confirmed the reproducibility of NAMPT, AKT, and BRAF lineage signals (Figure 6c).

**Figure 6.**
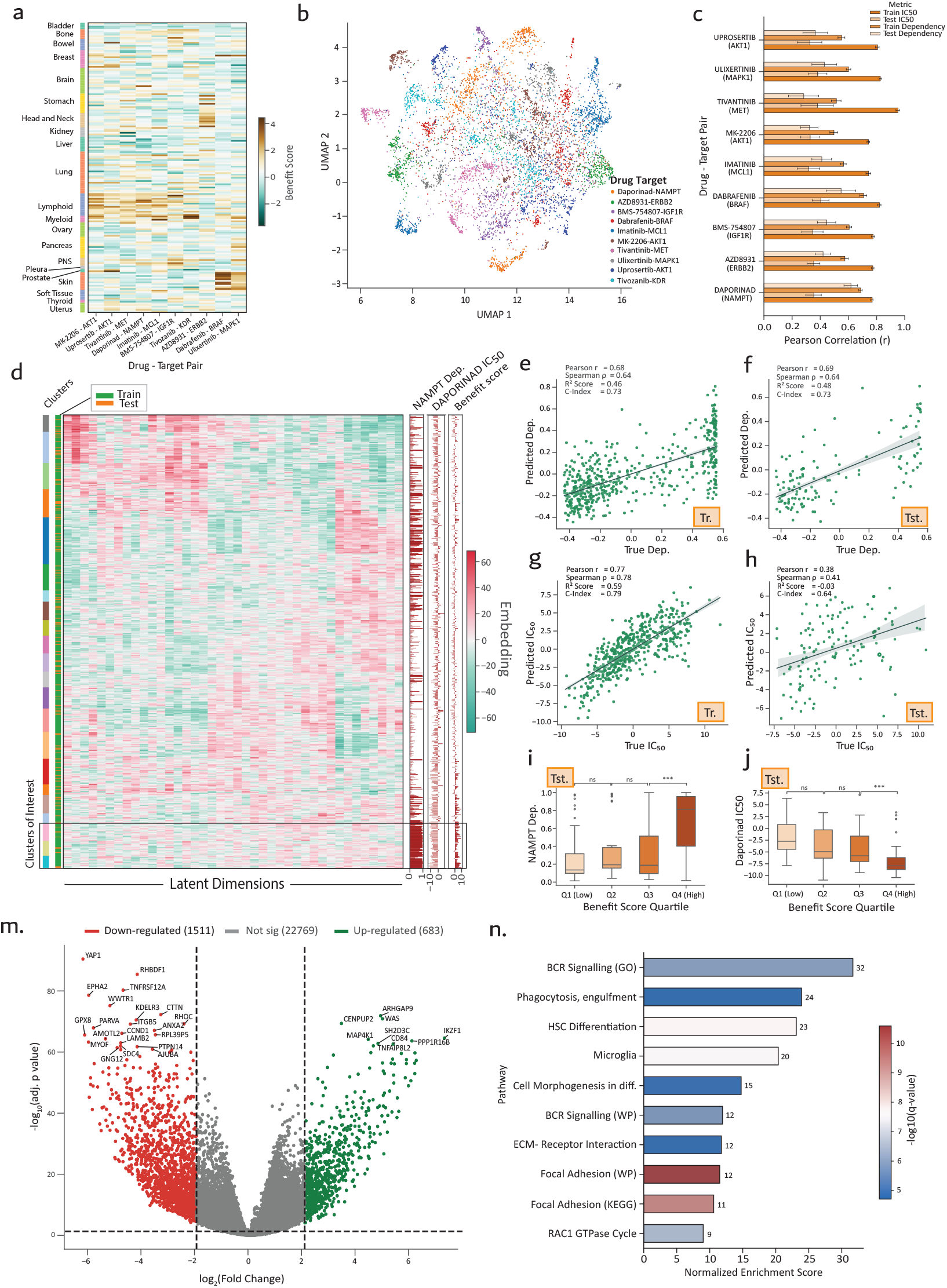
a-c**FORGE** model performances for other drug-target pairs, **d-n** Analysis of the NAPMT-Daporinad pair. **a**, Heatmap depicting the Benefit Score distribution of various drug-target pairs categorised according to the tissue sources. The data is sourced from DepMap. Many drug-target pairs had higher Benefit Scores for the lymphoid tumors. **b**, UMAP plot depicting the sample-level latent embeddings upon **FORGE** model training for various drug-target pairs. **c**, Barplot showing Pearson *r* correlation between model predictions and actual values. **d**,Heatmap of sample-level latent embeddings for NAMPT-Daporinad pair. Key clsuters with high dependency-benfitscore-low IC_50_ can be visualised **e-h**, Scatterplot and performance metrics for the NAMPT-Daporinad pair. **i-j**, Boxplots depicting the distributions of target dependency and drug IC_50_ among various groups based on Benefit Scores. **k**, Volcano plot depicting differential gene expression among high and low Benefit Score categories. **l**, Gene set enrichment analysis revealing top significantly enriched terms from the enrichr-KG.

For the NAMPT–Daporinad pair, model’s predictions had moderate to strong correlation for both dependency (Pearson’s *r* = 0.69) and IC_50_ (Pearson’s *r* = 0.38) (Figure 6e-h). Differential expression and enrichment analyses revealed that sensitive lines upregulated immune and metabolic “brake” genes (e.g., ARHGAP9 [60], TNFAIP8L2 [61], CD84 [62], IKZF1) [63], which suppress proliferative signaling or enforce immune checkpoint–like states. Conversely, downregulated genes included proliferative and migratory drivers such as YAP1 [64], WWTR1/TAZ, EPHA2 [65], CTTN [66], and CCND1, pointing to weakened Hippo signaling and adhesion programs (Figure 6m). This transcriptional profile—immune-regulatory, low-proliferation, and metabolically constrained—suggests that sensitive cells are unable to buffer the NAD^+^/ATP collapse induced by NAMPT inhibition. Enrichment highlighted blood and immune pathways, including focal adhesion, leukemia, and lymphoma signatures [67, 68], reinforcing the biological plausibility of NAMPT dependence in hematologic contexts [69] (Figure 6n).

**FORGE** generalizes to multiple drug–target contexts, reliably recovering lineage–drug associations aligned with known indications while also uncovering unexpected patterns, such as strong tivozanib (KDR) signals in myeloid lineages. These novel enrichments are best interpreted as potential repositioning hypotheses, suggesting lineage-specific contexts that warrant further preclinical or clinical investigation. By integrating Benefit Score stratification with differential expression and enrichment analysis, **FORGE** provides not only statistical robustness but also biologically grounded interpretability, positioning it as a framework for biomarker discovery and the rational exploration of new therapeutic opportunities.

## Discussion

Our study describes a novel implementation of joint-matrix factorization called **FORGE**, and its critical evaluation by validating on independent datasets. We also demonstrated the applicability of **FORGE** in understanding therapeutic responses of the EGFR-erlotinib pair, and in modeling multiple other drug-target pairs. Our implementation merits in using simple but powerful matrix factorization method to jointly learn two related biological variables - the gene’s dependency and drug’s IC_50_. Using these jointly learned embeddings, we derived a quantifiable score, the **Benefit Score**, which can be used in predicting prognosis for targeted therapies in cancer. Through this approach, we successfully bridged pharmacogenomics and functional genomics in a single framework that can be used for personalized therapy design. Our joint-model, in particular, identified key genetic signatures that can drive therapy responses. Our model formulation involved using numerous publicly available resources - the DepMap and the CREAMMIST databases, which provide high-quality cell line level influences of gene knockouts and the pharmacodynamics of numerous drugs either approved or under active research.

Classifying patients into **high** and **low** responders remains a holy grail in cancer research and facilitates the design of personalized therapies. Earlier methods predominantly relied on multiple neural network architectures, such as graph neural networks (CDRscan) [10] and autoencoders (DeepDR) [12] to predict drug responses using various DL architectures. These models often integrated various omics datasets, and even included drug structure information in training [70]. However, very few approaches utilized transcriptomic data [70], and deep learning-based models are generally compute-intensive and generate **black-box** predictions that rarely help in understanding the biological mechanisms driving model outputs. On the other hand, simple linear models have been shown to be effective in interpreting perturbation effects in single-cell data [71, 72].

**FORGE**’s validation using multiple independent datasets shows the model both stratifies samples meaningfully and offers interpretable mechanistic signals. In the DepMap-CREAMMIST cohort, **FORGE** identified compact clusters of sensitive cell lines (notably NSCLC and HNSCC) that align with established EGFR biology. The sparsity-inducer in **W** matrix concentrates influence on a small set of genes, producing gene-influence maps that reduce noise and are straightforward to interpret. Put simply, **FORGE** keeps a balance between accuracy and transparency: it achieves competitive prediction while giving clear, biologically sensible explanations for those predictions. We further demonstrated portability across biological contexts. Applying learned gene weights to external datasets (Tahoe-100M single-cell perturbations and PDX xenografts) retained signal: Benefit Scores separated malignant from normal cell lines and highlighted cells with larger transcriptomic reprogramming after treatment in Tahoe-100M, and in the PDX collection high Benefit Scores were associated with greater tumor regression. These cross-system validations show that **FORGE**’s embeddings and Benefit Score are not idiosyncratic to a single dataset but capture reproducible, treatment-relevant biology across cell lines, single-cell perturbations and *in vivo* models. We even demonstrated the Benefit Scores are not meaningful if an unrelated gene (A1CF) has been paired with a drug (see Supplementary Note 1).

Biologically, **FORGE** identifies sensitizers and resistance factors that are consistent with known EGFR mechanisms (for example, AREG and GRB7 as sensitizers; TNK1 and PPL as resistance-associated). Unlike a simple differential expression analysis, **FORGE** aligns gene effects across dependency and drug response simultaneously, so the prioritized genes are more likely to be mechanistic drivers of sensitivity or resistance rather than context-specific expression changes. **FORGE** also recovers lineage-specific vulnerabilities (e.g., NAMPT in hematologic contexts, BRAF in skin) and flags unexpected enrichments (such as strong KDR/Tivozanib signal in myeloid lines) that serve as testable repositioning hypotheses. In short, **FORGE** is both a discovery engine and a hypothesis generator for translational follow-up. Conceptually, the benefit of the joint embedding is straightforward: by forcing the same latent factors to explain both dependency and IC_50_, the model reduces task-specific noise and aligns pharmacologic readouts with functional genetic evidence. That alignment yields coherent molecular drivers that support clinically meaningful stratification strategies — i.e., groups defined by shared latent programs that map to pathway biology and likely treatment benefit.

Limitations remain. The linear factorization used in **FORGE** may miss highly nonlinear gene–drug interactions, and the current design requires training a model per drug–target pair (a trade-off we accept because per-target models are more useful for personalized therapy). External validations such as the PDX set were hampered by missing genes and small sample sizes, reducing statistical power. The Benefit Score depends on accurate baseline expression and can be affected by residual batch effects if not handled during the preprocessing steps. Finally, the present implementation targets single drugs or small drug sets; extending **FORGE** to polytherapy or to incorporate other omics or drug-structure features is a promising direction but raises the question of whether the core linear framework should be modified or the input matrix *G* simply augmented.

In summary, **FORGE** is a simple yet powerful framework that, to our knowledge, is the first tractable model to jointly link CRISPR-based gene dependency and IC_50_ drug response. It combines predictive performance with mechanistic interpretability, producing Benefit Scores that function as practical, transferable biomarkers and exposing both confirmatory biology and novel repositioning hypotheses. **FORGE** therefore offers a generalizable, mechanism-aware tool for biomarker discovery and the rational stratification of targeted therapies.

## Methods

### Overview of the datasets

Below we list down key datasets used in our study. The DepMap and the CREAMMIST datasets were utilized for model training, while model validation was performed using Tahoe-100M and PDX datasets.

#### DepMap datasets

We utilized the ‘Public 24Q4’ version of the DepMap (https://depmap.org/portal/) database, that contains gene expression (‘omicsExpressionRNASeQCGeneCountProfile.csv’) profiles of 53,970 genes among 1,690 cancer cell lines. Of these, 1,178 cell lines were subjected to CRISPR-Cas9-mediated knockouts targeting 17,917 genes. The effects of gene perturbation are estimated as gene dependency values (‘CRISPRGeneDependency.csv’) that range between 0 (**no effect**) and 1 (**highly dependent**).

#### CREAMMIST database

The CREAMMIST (https://creammist.mtms.dev/drug/select/) database contains log_10_-scaled half-maximal inhibitory concentrations (hereafter referred as IC_50_) of 1,132 drugs tested in 1,325 cancer cell lines belonging to 31 different cancer types. From this list of drugs, we included the drugs annotated in both OpenTargets (https://www.opentargets.org/) and TTD (https://idrblab.net/ttd/) databases, and have at least one known molecular target annotated in the CREAMMIST database, leaving a final count of 184 drugs for subsequent analysis.

#### Tahoe 100M scRNAseq

The Arc Institute (https://arcinstitute.org/news/arc-vevo) recently published a large-scale single-cell perturbation dataset comprising of 100M cells belonging to 49 cancer and one normal (HERT-hPNE) cell lines [73]. The atlas was derived through their Mosaic platform, and measured the effects of 1,100 small molecule perturbations at varying concentrations. Each plate (the batch variable) included a specific concentration of the perturbagen and DMSO (as a control), and were sequenced to measure the expression levels of 62,710 transcripts.

#### Patient-derived Xenografts (PDX)

This dataset is composed of baseline gene expression (FPKM) measuremnts of ∼1000 PDX samples belonging to six cancer types including the non-small cell lung cancer (NSCLC) [74]. The tumors are then treated using a variety of anti-cancer drugs, among which 25 samples were treated with erlotinib. Treatment responses were recorded as percentage change in tumor volumes (%TVChange) when compared with baseline (*t*=0) across different time points.

### Preprocessing

Preprocessing was carried out separately for training and each of the validation datasets (Supplementary Figure 1) and included voom normalization of raw gene counts, and subsequent scaling using Z-scores. The DepMap gene expression and dependency data, and the IC_50_ data from CREAMMIST databases together form the training datasets, while model validation was performed using the PDX and the Tahoe-100M datasets. We initially focused on EGFR-erlotinib pair for **FORGE** implementation, and the pre-processing steps described below were specific to this pair. We then expanded **FORGE** on other drug-target pairs using the same pre-processing schema.

We initially filtered the common cell lines in all the three training datasets, leaving a total count of 658 cell lines for downstream analysis. The cancer types and tissue sources for these cell lines are presented in the (Supplementary Figure 2a-b). The gene expression dataset is further filtered to include genes (N=24,963) present in enrichr-KG (https://maayanlab.cloud/enrichr-kg), a knowledge-graph based gene set enrichment tool. Subsequently, the gene expression was normalized using Limma’s Voom normalization with only the intercept in the design formula, and using default parameters. Both the gene dependency and drug IC_50_ values were scaled using their mean and represented the output variables for model training.

#### Feature selection and generating input matrices

From the above datasets, the data was split into train-test fractions in 80:20 ratio. Given the large number of input features, we identified top 200 genes (hereafter referred as highly-correlated genes or **HCGs**) with highest correlation with either of the output variables in the training data alone. This subset of gene expression values were then scaled through Z-transformations and constituted the training feature set **G**. The mean-centered gene dependency and drug IC_50_ values formed the dependent features - **D** and **I** respectively.

#### The PDX dataset

The original dataset contained FPKM values derived from bulk RNAseq and mapping the reads to the GRCh37 (along with mm9) genome build. We initially subsetted the data to include the 25 PDX models treated with erlotinib alone, and converted gene names into HGNC symbols using customized R script. Subsequently we converted the FPKM counts into pseudo-raw counts by using the original FPKM formula,

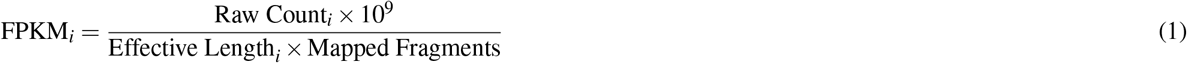

Solving for raw counts,

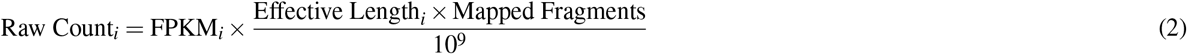

We then assumed a typical RNA-seq depth of approximately 35 *×* 10^6^ mapped fragments, and effective gene length as *L*_*i*_ − 199, where *L*_*i*_ is the full gene length and 199 bp approximates the average length of 75-100 bp paired-end sequencing fragments. Further simplification yields:

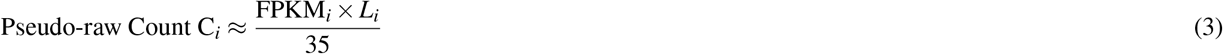

where FPKM_*i*_ and *L*_*i*_ refer to the gene’s FPKM value and effective gene length for gene *i*, respectively (for detailed derivation and an example, refer Supplementary file 2). We retrieved the gene lengths for the GRCh37 build using a customized python script, and transformed these pseudo-raw counts using Limma Voom normalization and Z-scaling. We used the percent change in tumor volume (%TVChange) as the main response variable, along with Best-average-response as implemented in the original publication [74], and binned the %TVChange values into durations spanning five days.

#### Tahoe 100M dataset

From the Tahoe-100M dataset, we selected plates 7, 8 and 9 as they constitute the samples treated with erlotinib at three different concentrations - 0.05 nM (**low**), 0.5 nM (**medium**) and 5.0 nM (**high**). We extracted the drug and DMSO treated cells from these plates into separate AnnData files, and aggregated the cell-line-specific raw counts into pseudo-bulk count matrices. For computing the Benefit Scores, we used the pseudo-bulk counts from plate 9’s DMSO-treated cell-lines that were then fed to the data normalization pipeline that included Limma’s Voom normalization and Z-transformations. The drug-treated cell-lines were normalized separately and both these normalizations had only the intercept in the design formula.

### FORGE

The **FORGE** model is a joint matrix factorization framework designed to integrate gene expression profiles with functional screening data. It takes baseline gene expression data **G** ∈ ℝ^*n×p*^ as input, where *n* and *p* represent number of samples and genes respectively. The model learns a shared sample-level embedding that captures predictive features relevant to both gene dependency (**D**) and drug response (measured by IC_50_, denoted **I**).

The model approximates these two response matrices as:

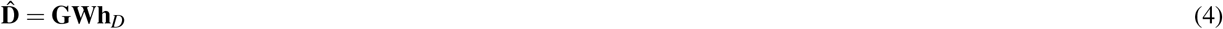

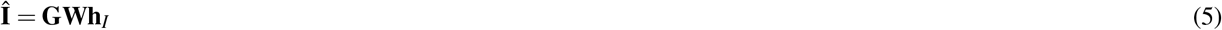

Here, **W** ∈ ℝ^*p×k*^ represents the gene expression embeddings in a latent space of dimension *k* (a tunable hyperparameter), while **h**_**D**_ ∈ ℝ^*k×*1^ and **h**_**I**_ ∈ ℝ^*k×*1^ denote the projection vectors for gene dependency and drug response, respectively.

Model training is performed by minimizing a composite loss function that balances reconstruction accuracy and sparsity:

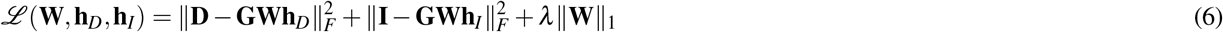

The regularization parameter *λ* was set to a default value of 10^−3^. The gradient of this objective with respect to **W** is given by:

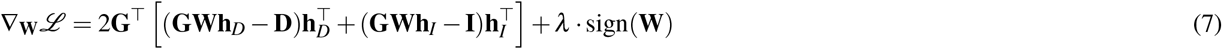

The parameter matrix **W** is updated iteratively using gradient descent:

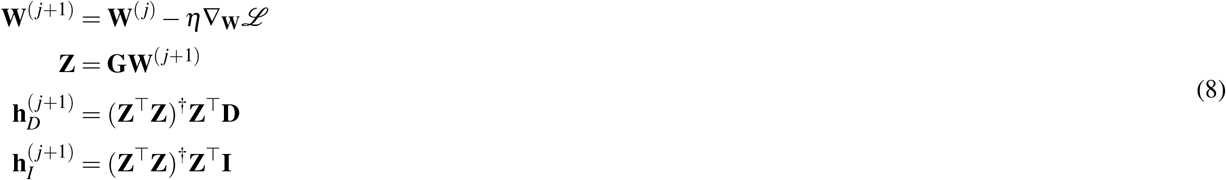

where *η* is the learning rate (another hyperparameter with the default value set at 0.01), and † denotes the Moore–Penrose pseudo-inverse. The inclusion of an *L*_1_ penalty on **W** encourages sparsity in the learned representations. To determine the optimal latent dimensionality *k*, we performed a grid search across values ranging from 10 to 150. For each value of *k*, five independent runs were conducted with randomly initialized **W**. Performance was evaluated by computing the Pearson *r* correlation between observed (**D**,**I**) and predicted (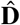 and ***Î***) values across both training and held-out test sets. The *k* value that maximized average predictive accuracy was selected for downstream analyses.

### Benefit Score

The **benefit score** is a quantitative estimate per sample combining sample-specific baseline expression profile with the learned embeddings from the **FORGE** model. For making the computations tractable, we initially computed gene-wise **influence scores** as:

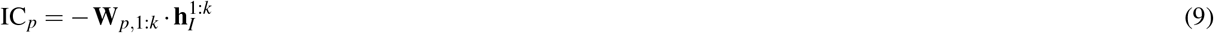

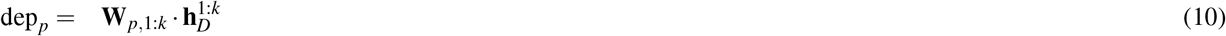

where **W**_*p*,1:*k*_ denotes the weight vector for gene, 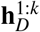 and 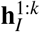 denote the learned embedding vectors for the dependency and IC_50_ tasks, respectively. The negative sign in IC_*p*_ reflects the inverse relationship between IC_50_ and benefit. Finally, we aggregated gene’s influence scores as:

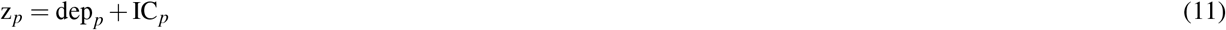

This formulation is equivalent to computing:

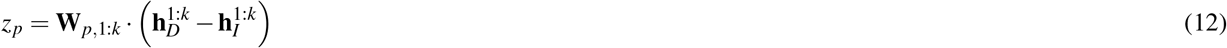

Finally, the benefit score for a sample *i* was estimated as:

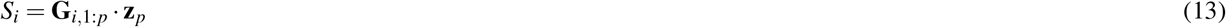

For understanding the model’s learned representations, we plotted the correlation between dependency and IC_50_ influence scores. Additionally, we used the median of the scores in each dataset to segregate the samples into two groups - **high-score** and **low-score** for group-wise comparisons of gene expression changes and response variables.

### FORGE: Individual model

In the individual modeling framework, we considered a single prediction task at a time, either gene dependency (**D**) or drug response (**I**). Let **G** ∈ ℝ^*n×p*^ denote the gene expression matrix, where *n* is the number of samples (cell lines or PDX models) and *p* is the number of genes. The aim is to learn a mapping from the high-dimensional input space (*p* genes) into a latent embedding that explains either dependency profiles or drug sensitivity values.

#### Model definition

For the gene dependency prediction task, the model takes the form

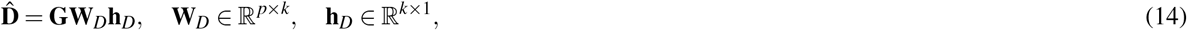

where *k* is the latent dimensionality, **W**_*D*_ represents the projection of gene expression features into the latent space, and **h**_*D*_ denotes the task-specific weights that combine these embeddings into the prediction 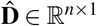.

Similarly, for drug response prediction (IC_50_), the model is defined as

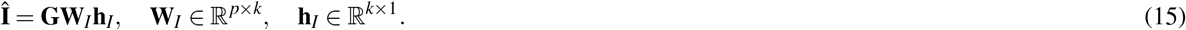

#### Loss function

The optimization objective for each individual model balances fidelity to the observed data with a sparsity-inducing regularizer on the weight matrix:

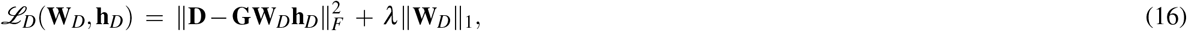

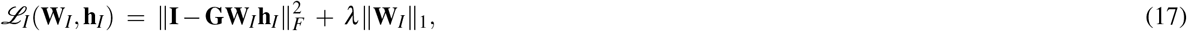

where ∥·∥_*F*_ denotes the Frobenius norm, and ∥·∥_1_ denotes the element-wise 𝓁_1_ norm. The first term enforces reconstruction accuracy (minimizing the squared error between observed and predicted values), while the second term encourages sparsity in the gene-to-latent projection matrix, thereby identifying a compact set of informative features. The regularization coefficient *λ >* 0 controls the trade-off between reconstruction accuracy and sparsity.

#### Interpretation of parameters

**W**_*D*_ and **W**_*I*_ capture the mapping from gene expression profiles to latent factors relevant to each task. - **h**_*D*_ and **h**_*I*_ represent task-specific linear coefficients that weight the latent factors for predicting dependency or IC_50_, respectively. - The sparsity regularizer *λ*∥**W**∥_1_ ensures interpretability by shrinking irrelevant gene weights toward zero.

#### Optimization

Each loss function (16) and (17) was minimized independently using stochastic gradient descent over a maximum of 500 epochs. Five independent runs with different random initializations were performed to account for stochastic variability. The latent dimensionality *k* was fixed to the value optimized in the joint model setting.

#### Evaluation metrics

The performance of the trained models was quantified by computing Spearman’s rank correlation coefficient (*ρ*) to measure monotonic agreement, and mean absolute error (MAE) to assess average predictive deviation. These metrics were averaged across the five replicate runs, and the resulting distributions were reported for each learning task.

### Multi-drug FORGE

To accommodate multiple drugs targeting the same molecular target, we extended the **FORGE** framework into a multi-output setting. In this formulation, the response variable **I** is no longer a vector of IC_50_ values, but a matrix capturing responses across multiple drugs.

#### Model definition

Let **G** ∈ ℝ^*n×p*^ denote the gene expression matrix with *n* samples and *p* genes, as in the individual model. For *d* drugs targeting the same gene, the drug response matrix is

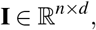

where each column corresponds to the IC_50_ profile of a specific drug across all *n* samples.

The prediction model is generalized to

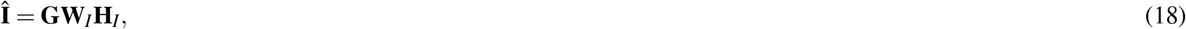

where

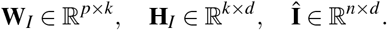

Here, **W**_*I*_ projects the high-dimensional gene expression features into a *k*-dimensional latent space, and **H**_*I*_ captures task-specific (drug-specific) coefficients that map the latent representations to *d* parallel prediction outputs. The prediction for dependency remains the same and is generalized as:

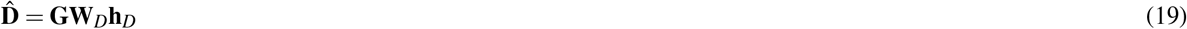

#### Loss function

The learning objective for the multi-drug model is defined as:

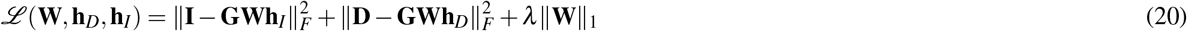

where ∥·∥_*F*_ denotes the Frobenius norm, and ∥·∥_1_ is the element-wise 𝓁_1_ norm of **W**_*I*_. The first term enforces accurate reconstruction of all *d* drug response profiles simultaneously, while the dependency reconstruction remained the same. Similar to that of individual and joint **FORGE** models, the third term induces sparsity in **W** to encourage selection of informative genes.

#### Interpretation

**W**_*I*_ represents gene-to-latent feature mappings that are shared across all drugs targeting the same gene. - **H**_*I*_ is a task-specific coefficient matrix, where each column corresponds to the weights associated with one drug, thereby disentangling common vs. drug-specific signals. - **I** captures the observed variability in drug sensitivity across multiple drugs, while **Î**represents the corresponding model predictions.

#### Application to EGFR inhibitors

To illustrate this framework, we considered two EGFR inhibitors, Erlotinib and Gefitinib, which exhibit substantial heterogeneity in IC_50_ distributions across cell lines. In this case, **I** is a two-column matrix:

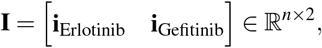

where **i**_drug_ represents the IC_50_ values for a given drug across *n* samples. The model was trained using the same optimization scheme as in the single-task setting, with the latent dimensionality *k* fixed to the value optimized in the joint framework and a maximum of 500 epochs. Five replicate runs with different random initializations were conducted.

#### Evaluation

Model performance was evaluated on the DepMap dataset by computing (i) Spearman’s rank correlation coefficient (*ρ*) between predicted and observed IC_50_ values for each drug separately, and (ii) mean absolute error (MAE) across the *n×d* prediction matrix. This allowed us to quantify the ability of the shared latent structure to jointly explain drug responses that target the same pathway.

### Tuning model hyper-parameters

For the **FORGE** model, one of the key hyperparameters is the number of latent dimensions, *k*. We tested different values for *k* ranging between 10 and 150 with a step-size of 10. For each such iteration, we used five bootstrap replicates (labeled b0 to b4), and each iteration used a different random initialization for **W**. Subsequently, we identified the best *k* by considering the value that resulted in highest correlation between predicted and actual values for train and test datasets. The other hyperparameters—*λ* (default value 1e-3), *η* (default value 1e-3), and the number of epochs (500)—were chosen arbitrarily.

### FORGE training complexity

The computational cost of training *FORGE* mainly comes from the matrix multiplications involved in computing gradients for the shared weight matrix *W*. Each gradient evaluation scales as 𝒪(*npk*), where *n* is the number of cell lines, *p* is the number of genes, and *k* is the latent dimension. For the task-specific heads (*h*_*D*_ and *h*_*I*_), the updates can be computed in closed form and require 𝒪(*nk*^2^ + *k*^3^) operations. Since *k* is relatively small (e.g., *k* = 40), this cost is negligible compared to the gradient updates. If training involves *T* outer alternating iterations (typically 100), and each update of *W* uses *S* proximal gradient steps (typically 10), then the overall computational complexity of the algorithm is

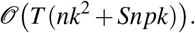

### FORGE performance evaluations

For evaluating the model’s performance, we used a variety of metrics - the Pearson *r*, Spearman *ρ*, and *R*^2^ for estimating correlations and goodness of fit between actual and predicted values. We also used the concordance index, a metric that quantifies the concordance between model predictions and the actual values among all such comparisons.

### Response strength

For the Tahoe 100M dataset, we computed **response strengths** alongside benefit scores as the key response variable. This metric captures the magnitude of transcriptomic shifts induced by drug treatment. Response strengths were subsequently compared across the three drug concentrations among samples stratified by benefit score categories.

Let **x**_drug_ = [*x*_1_, *x*_2_, …, *x*_*n*_] and **x**_control_ = [*c*_1_, *c*_2_, …, *c*_*n*_] denote the normalized expression vectors for a sample under drug and control conditions respectively. Then:

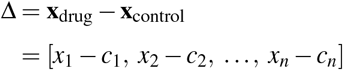

The response strength is defined as the Euclidean norm of the difference vector Δ:

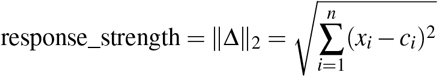

### Differential gene expression analysis

For the differential expression analyses for the DepMap and the Tahoe 100M datasets, the samples were categorized into two groups (**high-score** and **low-score**) using the median Benefit Score of each dataset as cutoffs. This variable was the used as the main contrast, and we included cancer type or tissue type (whenever applicable) as one of the co-variates. For the DepMap dataset, differential expression analyses were carried out using Limma’s ebayesR using the existing normalized counts, while for the Tahoe-100M pseudo-bulk counts, we used DESeq2 and specified the counts as ‘poscounts’ in the *estimateSizeFactors()* step accounting for sparsity observed in the single-cell transcriptomes. The top 200 differentially expressed genes (DEGs) were identified based on abs(log_2_Fold change) > 1 and adjusted P < 0.05.

Alternately, for the DepMap dataset with 1034 cell lines that have both EGFR dependency and gene expression data, we performed additional DEG analysis. Based on the learned gene expression embeddings (**Z = G W**), we clustered the cell lines using hierarchical clustering using pairwise Euclidean distances and Ward’s method which minimizes within cluster variance. Subsequently, we resolved clusters using *cutreeHybrid* from the ***dynamicTreeCut*** library in Python by setting a *minClusterSize* of 13 (minimum number of samples per cluster) and *deepSplit* of 4 (maximize the cluster split, ranges from 1 to 5). From the resulting clusters, we aggregated samples from top three clusters with high benefit scores and EGFR dependency and low erlotinib IC_50_, and compared differential expression profiles with other samples.

The DEGs from each analysis were then used as separate input lists for the gene set enrichment analysis (GSEA) using enrichr-KG (https://maayanlab.cloud/enrichr-kg). We selected five databases to infer the gene functions in metabolism and disease - the GO Biological Process v2021, the WikiPathways v2021, the KEGG Pathway v2021 (for human), the Reactome pathways v2022, and the DisGeNET databases. We limited the results to gene sets of sizes ranging between 10 and 500 genes, at least 10 genes from the input list were mapped to each set, and a gene to be mapped to at least three different gene sets. All the enriched terms were filtered based on FDR q-value < 0.05, and validated our findings through extensive literature search.

### Statistical analysis

We used the Benefit Scores for comparing therapy outcome variables by categorizing samples into *high* and *low* scoring samples. The final outcome variables used for assessing Benefit Scores are dataset-specific – response strengths for Tahoe-100M dataset and %TVChange in PDX datasets. For all the above analyses, *P*<0.05 was considered as the level of statistical significance, and the following hypotheses were tested using one-sided Kolmogrov-Smirnov or Mann-Whitney tests:

H_0_ (the null hypothesis): There is no difference of response variable among the two groups.

H_1_ (the alternate hypothesis): The response strength is lower or the %TVChange was higher among the samples with low Benefit Scores (based upon the dataset tested).

Statistical significance of performance differences between multi-task and single-task models for multiple drug-target pairs were assessed using a paired *t*-test on correlation between **DependencyEffect** and **IC**50**Effect**.

### Software versions, code and data availability

Analyses were performed using R (v4.4.3) and Python (v3.10.12). The Limma package (v3.58.1) and ScanPy (v1.11.1) were applied where appropriate. Publicly available datasets from the DepMap[6] and CREAMMIST [7] resources were used in this study. Patient-derived xenograft (PDX) data were obtained from a previously published study [74], and the Tahoe-100M dataset was accessed through the Arc Institute [73]. The full code, including the main **FORGE** pipeline, is available at https://github.com/sreerampeela/FORGE.

## Acknowledgements

NB acknowledges research fellowship awarded by the Department of Biotechnology (DBT) – India (file no: DBTHRDPMU/JRF/BET-24/I/2024-25/306). DS acknowledges funding from DBT (IC-12044(12)/4/2022-ICD-DBT) and the Indian Council of Medical Research (ICMR) – India (EMDR/IG/16/2024-01-01321), and Science and Engineering Research Board (SERB) – India (CRG/2022/007706).

## Author contributions statement

D.S. conceived the idea; N.B. and S.C.M.P. carried out the data analysis and compiled the manuscript; A.H., B.M., R.G. and S.S. helped in critical evaluation of results and data visualization; S.S., S.K. and S.P. reviewed the manuscript; G.A., S.D., A.M. and D.S. reviewed the manuscript and provided critical insights into the results. A.M. provided the mathematical proof of the FORGE model. N.B. and S.C.M.P. contributed equally to this work.

## References

1. World Health Organization. Cancer https://www.who.int/health-topics/cancer. Accessed: 23 August 2025.

2. Allison, K. H. & Sledge, G. W. Heterogeneity and cancer. Oncology 28, 772–772 (2014).

3. Larrimore, K. E. & Rancati, G. The conditional nature of gene essentiality. Current Opinion in Genetics & Development 58, 55–61 (2019).

4. Abdullah, S. E., Haigentz Jr, M. & Piperdi, B. Dermatologic toxicities from monoclonal antibodies and tyrosine kinase inhibitors against EGFR: pathophysiology and management. Chemotherapy research and practice 2012, 351210 (2012).

5. Li, Y. et al. Mechanism of lethal skin toxicities induced by epidermal growth factor receptor inhibitors and related treatment strategies. Frontiers in Oncology 12, 804212 (2022).

6. Arafeh, R., Shibue, T., Dempster, J. M., Hahn, W. C. & Vazquez, F. The present and future of the Cancer Dependency Map. Nature Reviews Cancer 25, 59–73 (2025).

7. Yingtaweesittikul, H. et al. CREAMMIST: an integrative probabilistic database for cancer drug response prediction. Nucleic Acids Research 51, D1242–D1248 (2023).

8. Nusinow, D. P. et al. Quantitative proteomics of the cancer cell line encyclopedia. Cell 180, 387–402 (2020).

9. Ganini, C. et al. Global mapping of cancers: The Cancer Genome Atlas and beyond. Molecular oncology 15, 2823–2840 (2021).

10. Chang, Y. et al. Cancer drug response profile scan (CDRscan): a deep learning model that predicts drug effectiveness from cancer genomic signature. Scientific reports 8, 8857 (2018).

11. Jiang, L. et al. DeepTTA: a transformer-based model for predicting cancer drug response. Briefings in bioinformatics 23 (2022).

12. Jiang, Z. & Li, P. DeepDR: a deep learning library for drug response prediction. Bioinformatics 40, btae688 (2024).

13. Chawla, S. et al. Gene expression based inference of cancer drug sensitivity. Nature communications 13, 5680 (2022).

14. Chawla, S. et al. UniPath: a uniform approach for pathway and gene-set based analysis of heterogeneity in single-cell epigenome and transcriptome profiles. Nucleic acids research 49, e13–e13 (2021).

15. Zhang, X., Xiao, W. & Xiao, W. DeepHE: Accurately predicting human essential genes based on deep learning. PLOS Computational Biology 16, e1008229 (2020).

16. Kuang, S., Wei, Y. & Wang, L. Expression-based prediction of human essential genes and candidate lncRNAs in cancer cells. Bioinformatics 37, 396–403 (2021).

17. Baker, R. E., Pena, J.-M., Jayamohan, J. & Jérusalem, A. Mechanistic models versus machine learning, a fight worth fighting for the biological community? Biology letters 14, 20170660 (2018).

18. Angermueller, C., Pärnamaa, T., Parts, L. & Stegle, O. Deep learning for computational biology. Molecular systems biology 12, 878 (2016).

19. Lee, B. D. et al. Ten quick tips for deep learning in biology. PLoS computational biology 18, e1009803 (2022).

20. Noordijk, B. et al. The rise of scientific machine learning: a perspective on combining mechanistic modelling with machine learning for systems biology. Frontiers in Systems Biology 4, 1407994 (2024).

21. Ovchinnikova, K., Born, J., Chouvardas, P., Rapsomaniki, M. & Kruithof-de Julio, M. Overcoming limitations in current measures of drug response may enable AI-driven precision oncology. NPJ precision oncology 8, 95 (2024).

22. Baião, A. R. et al. A technical review of multi-omics data integration methods: from classical statistical to deep generative approaches. Briefings in Bioinformatics 26, bbaf355 (2025).

23. Wang, L., Li, X., Zhang, L. & Gao, Q. Improved anticancer drug response prediction in cell lines using matrix factorization with similarity regularization. BMC cancer 17, 513 (2017).

24. Guan, N.-N. et al. Anticancer drug response prediction in cell lines using weighted graph regularized matrix factorization. Molecular Therapy Nucleic Acids 17, 164–174 (2019).

25. Emdadi, A. & Eslahchi, C. Dsplmf: a method for cancer drug sensitivity prediction using a novel regularization approach in logistic matrix factorization. Frontiers in genetics 11, 75 (2020).

26. Adam, G. et al. Machine learning approaches to drug response prediction: challenges and recent progress. NPJ precision oncology 4, 19 (2020).

27. Yang, W. et al. Genomics of Drug Sensitivity in Cancer (GDSC): a resource for therapeutic biomarker discovery in cancer cells. Nucleic acids research 41, D955–D961 (2012).

28. Seashore-Ludlow, B. et al. Harnessing connectivity in a large-scale small-molecule sensitivity dataset. Cancer discovery 5, 1210–1223 (2015).

29. Tsherniak, A. et al. Defining a cancer dependency map. Cell 170, 564–576 (2017).

30. Blackhall, F. H., Rehman, S. & Thatcher, N. Erlotinib in non-small cell lung cancer: a review. Expert opinion on pharmacotherapy 6, 995–1002 (2005).

31. Soulieres, D. et al. Multicenter phase II study of erlotinib, an oral epidermal growth factor receptor tyrosine kinase inhibitor, in patients with recurrent or metastatic squamous cell cancer of the head and neck. Journal of clinical oncology 22, 77–85 (2004).

32. Way, G. P., Zietz, M., Rubinetti, V., Himmelstein, D. S. & Greene, C. S. Compressing gene expression data using multiple latent space dimensionalities learns complementary biological representations. Genome biology 21, 109 (2020).

33. Zhao, S. et al. Efficacy and tolerability of erlotinib 100 mg/d vs. gefitinib 250 mg/d in EGFR-mutated advanced non-small cell lung cancer (E100VG250): an open-label, randomized, phase 2 study. Frontiers in Oncology 10, 587849 (2020).

34. Bronte, G. et al. Are erlotinib and gefitinib interchangeable, opposite or complementary for non-small cell lung cancer treatment? Biological, pharmacological and clinical aspects. Critical reviews in oncology/hematology 89, 300–313 (2014).

35. Fan, W.-C. et al. Different efficacies of erlotinib and gefitinib in taiwanese patients with advanced non-small cell lung cancer: a retrospective multicenter study. Journal of Thoracic Oncology 6, 148–155 (2011).

36. Jang, T.-H. et al. MicroRNA-485-5p targets keratin 17 to regulate oral cancer stemness and chemoresistance via the integrin/FAK/Src/ERK/β -catenin pathway. Journal of biomedical science 29, 42 (2022).

37. Quarta, S. et al. SERPINB3 induces epithelial–mesenchymal transition. The Journal of Pathology: A Journal of the Pathological Society of Great Britain and Ireland 221, 343–356 (2010).

38. Koese, M. et al. Annexin A6 is a scaffold for PKCα to promote EGFR inactivation. Oncogene 32, 2858–2872 (2013).

39. Tran, Q. T. et al. EGFR regulation of epidermal barrier function. Physiological genomics 44, 455–469 (2012).

40. Sutter, C. H. et al. EGF receptor signaling blocks aryl hydrocarbon receptor-mediated transcription and cell differentiation in human epidermal keratinocytes. Proceedings of the National Academy of Sciences 106, 4266–4271 (2009).

41. Murillas, R. et al. Expression of a dominant negative mutant of epidermal growth factor receptor in the epidermis of transgenic mice elicits striking alterations in hair follicle development and skin structure. The EMBO journal 14, 5216–5223 (1995).

42. Wen, Y. et al. MAPK1E322K mutation increases head and neck squamous cell carcinoma sensitivity to erlotinib through enhanced secretion of amphiregulin. Oncotarget 7, 23300 (2016).

43. Chu, P.-Y., Li, T.-K., Ding, S.-T.Lai, I.-R. & Shen, T.-L. EGF-induced Grb7 recruits and promotes Ras activity essential for the tumorigenicity of Sk-Br3 breast cancer cells. Journal of Biological Chemistry 285, 29279–29285 (2010).

44. Bill, A. et al. ANO1/TMEM16A interacts with EGFR and correlates with sensitivity to EGFR-targeting therapy in head and neck cancer. Oncotarget 6, 9173 (2015).

45. Huang, R., Li, L., Wang, Z. & Shen, K. A systemic pan-cancer analysis of MPZL3 as a potential prognostic biomarker and its correlation with immune infiltration and drug sensitivity in breast cancer. Frontiers in Oncology 12, 901728 (2022).

46. Henderson, M. C. et al. High-throughput RNAi screening identifies a role for TNK1 in growth and survival of pancreatic cancer cells. Molecular cancer research 9, 724–732 (2011).

47. Lee, H. M. et al. The 4717C> G polymorphism in periplakin modulates sensitivity to EGFR inhibitors. Scientific Reports 9, 2357 (2019).

48. Sigismund, S. et al. Clathrin-mediated internalization is essential for sustained EGFR signaling but dispensable for degradation. Developmental cell 15, 209–219 (2008).

49. Grandal, M. V. & Madshus, I. H. Epidermal growth factor receptor and cancer: control of oncogenic signalling by endocytosis. Journal of cellular and molecular medicine 12, 1527–1534 (2008).

50. Jo, U. et al. EGFR endocytosis is a novel therapeutic target in lung cancer with wild-type EGFR. Oncotarget 5, 1265 (2014).

51. Joly-Tonetti, N., Ondet, T., Monshouwer, M. & Stamatas, G. N. EGFR inhibitors switch keratinocytes from a proliferative to a differentiative phenotype affecting epidermal development and barrier function. BMC cancer 21, 5 (2021).

52. Badger, B. L. Why deep learning generalizes. arXiv preprint 2211.09639 (2022).

53. Zhang, J. et al. Docetaxel resistance-derived LINC01085 contributes to the immunotherapy of hormone-independent prostate cancer by activating the STING/MAVS signaling pathway. Cancer Letters 545, 215829 (2022).

54. Gentilcore, G. et al. Effect of dabrafenib on melanoma cell lines harbouring the BRAF V600D/R mutations. BMC cancer 13, 17 (2013).

55. Oki, Y. et al. Phase II study of an AKT inhibitor MK2206 in patients with relapsed or refractory lymphoma. British journal of haematology 171, 463–470 (2015).

56. Chen, E. C. et al. Targeting MET and FGFR in relapsed or refractory acute myeloid leukemia: preclinical and clinical findings, and signal transduction correlates. Clinical Cancer Research 29, 878–887 (2023).

57. Mitchell, S. R. et al. Selective targeting of NAMPT by KPT-9274 in acute myeloid leukemia. Blood advances 3, 242–255 (2019).

58. Padro, T. et al. Overexpression of vascular endothelial growth factor (VEGF) and its cellular receptor KDR (VEGFR-2) in the bone marrow of patients with acute myeloid leukemia. Leukemia 16, 1302–1310 (2002).

59. Zhang, Y. et al. VEGFR2 activity on myeloid cells mediates immune suppression in the tumor microenvironment. JCI insight 6, e150735 (2021).

60. Song, W. et al. Rho GTPase activating protein 9 (ARHGAP9) in human cancers. Recent Patents on Anti-Cancer Drug Discovery 17, 55–65 (2022).

61. Li, T. et al. Genome-wide analysis reveals TNFAIP8L2 as an immune checkpoint regulator of inflammation and metabolism. Molecular immunology 99, 154–162 (2018).

62. Mangiola, S. et al. Circulating immune cells exhibit distinct traits linked to metastatic burden in breast cancer. Breast Cancer Research 27, 73 (2025).

63. Chen, J. C., Perez-Lorenzo, R., Saenger, Y. M., Drake, C. G. & Christiano, A. M. IKZF1 enhances immune infiltrate recruitment in solid tumors and susceptibility to immunotherapy. Cell Systems 7, 92–103 (2018).

64. Wang, W. et al. AMPK modulates Hippo pathway activity to regulate energy homeostasis. Nature cell biology 17, 490–499 (2015).

65. Nasreen, N., Mohammed, K. A. & Antony, V. B. Silencing the receptor EphA2 suppresses the growth and haptotaxis of malignant mesothelioma cells. Cancer 107, 2425–2435 (2006).

66. Jing, X. et al. Cortactin promotes cell migration and invasion through upregulation of the dedicator of cytokinesis 1 expression in human colorectal cancer. Oncology Reports 36, 1946–1952 (2016).

67. Burley, T. A. et al. Elucidation of focal adhesion kinase as a modulator of migration and invasion and as a potential therapeutic target in chronic lymphocytic leukemia. Cancers 14, 1600 (2022).

68. Jones, C. L. et al. Nicotinamide metabolism mediates resistance to venetoclax in relapsed acute myeloid leukemia stem cells. Cell stem cell 27, 748–764 (2020).

69. Kozako, T. et al. Induction of apoptosis and autophagy by nampt inhibition in adult T-Cell leukemia/lymphoma and leukemic cell lines. Blood 128, 2327 (2016).

70. Abbasi, M., Carvalho, F. G., Ribeiro, B. & Arrais, J. P. Predicting drug activity against cancer through genomic profiles and SMILES. Artificial Intelligence in Medicine 150, 102820 (2024).

71. Ahlmann-Eltze, C., Huber, W. & Anders, S. Deep-learning-based gene perturbation effect prediction does not yet outperform simple linear baselines. Nature Methods, 1–5 (2025).

72. Kulm, S., Kofman, L., Mezey, J. & Elemento, O. Simple linear cancer risk prediction models with novel features outperform complex approaches. JCO Clinical Cancer Informatics 6, e2100166 (2022).

73. Zhang, J. et al. Tahoe-100M: A Giga-Scale Single-Cell Perturbation Atlas for Context-Dependent Gene Function and Cellular Modeling. bioRxiv. https://www.biorxiv.org/content/early/2025/02/24/2025.02.20.639398(2025).

74. Gao, H. et al. High-throughput screening using patient-derived tumor xenografts to predict clinical trial drug response. Nature medicine 21, 1318–1325 (2015).

