## Supplementary Material for "Mechanism-aware inference of response to targeted cancer therapies"

#### Supplementary Notes

##### Correlation of Gene-Level Effects: Coupled vs. Separate Formulations

We now formally establish that the proposed coupled formulation (Eq. (6)) yields more correlated gene-level effect vectors $Wh_D$ and $Wh_I$ than the separate formulations (Eqs. (16) and (17)).

In the context of analyzing the geometric relationship between the latent projections of gene dependency and drug response, we omit the regularization term in the loss function for simplicity. When the regularization parameter $\lambda$ is sufficiently small, its influence is dominated by the data-fitting terms, and the key properties of the shared subspace projection can be examined without loss of generality.

Let $Wh_D$ and $Wh_I$ denote the gene-level effect vectors obtained from the coupled formulation (Eq. (6)), and let $W_Dh_D$, $W_Ih_I$ denote those from the separate formulations (Eqs. (16)-(17)). Then:

1. For latent dimension $k = 1$, the coupled formulation yields

   $$corr(Wh_D, Wh_I) = \pm 1$$

   while the separate formulations generally yield strictly smaller correlation.

2. For latent dimension $k > 1$, the coupled formulation selects a common subspace $\mathcal{U}$ maximizing the joint projected signal from both $D$ and $I$, thereby increasing the cosine similarity between $Wh_D$ and $Wh_I$ relative to the independent subspaces $\mathcal{U}_D$, $\mathcal{U}_I$ of the separate models.

Let $Z = GW$. Eliminating $h_D$ and $h_I$ via Eq. (8) gives

$$h_D = (Z^\top Z)^\dagger Z^\top D, \quad h_I = (Z^\top Z)^\dagger Z^\top I$$

and hence

$$Wh_D = A_W D, \quad Wh_I = A_W I, \quad A_W := W(Z^\top Z)^\dagger Z^\top$$

**Case** $k = 1$ Here $W = w \in R^{p\times 1}$, $Z = Gw$. Eq. (8) simplifies to

$$h_D = \frac{Z^\top D}{Z^\top Z}$$

so that

$$h_I = \frac{Z^\top I}{Z^\top Z}$$

$$Wh_D = w\frac{Z^\top D}{Z^\top Z}, \quad Wh_I = w\frac{Z^\top I}{Z^\top Z}$$

Both are scalar multiples of the same direction $w$, hence colinear. Thus $corr(Wh_D, Wh_I) = \pm 1$. In contrast, the separate formulations yield distinct directions $w_D$, $w_I$ and therefore strictly smaller correlation unless they coincide by chance.

**Case** $k > 1$ For general $k$, note that $Wh_D = P_{\mathcal{U}}D$, $Wh_I = P_{\mathcal{U}}I$ where $\mathcal{U} = col(GW)$ and $P_{\mathcal{U}}$ is the projection onto $\mathcal{U}$. Substituting the optimal $h_D$, $h_I$ into the joint loss (Eq. (6)) gives

$$\min_{\mathcal{U}} \|(I - P_{\mathcal{U}})D\|_2^2 + \|(I - P_{\mathcal{U}})I\|_2^2,$$

equivalently maximizing the total projected energy $\|P_{\mathcal{U}}D\|^2 + \|P_{\mathcal{U}}I\|^2$. If $D$ and $I$ share signal (nonzero cross-covariance), the optimizer aligns $\mathcal{U}$ with shared directions, increasing

$$\langle P_{\mathcal{U}}D, P_{\mathcal{U}}I\rangle = D^\top P_{\mathcal{U}}I$$

and thus the cosine similarity

$$\cos\angle(Wh_D, Wh_I) = \frac{\langle P_{\mathcal{U}}D, P_{\mathcal{U}}I\rangle}{\|P_{\mathcal{U}}D\|\|P_{\mathcal{U}}I\|}.$$

In the separate case, distinct subspaces $\mathcal{U}_D, \mathcal{U}_I$ are chosen independently, so no such alignment is enforced, and correlation is weaker.

From the above derivation, the maximal value of $\theta$ is obtained when $P_U\mathbf{D}$ and $P_U\mathbf{I}$ are in the same direction, which can occur if and only if they share the same subspace. On the other hand, if $\mathbf{h}_D$ and $\mathbf{h}_I$ are optimized separately, the subspaces of $\mathbf{W}_D$ and $\mathbf{W}_I$ are different, leading to $\theta \neq 0$.

This proves the claim.

#### Selection of Latent Dimension $k = 40$

The number of latent dimensions ($k$) is a key hyperparameter in the *FORGE* framework, as it controls the balance between model expressiveness and generalizability. To systematically determine an appropriate value, we performed five independent bootstrap iterations across a range of latent dimensions (from $k = 10$ to $k = 150$) and evaluated model performance on both training and test sets.

Two criteria guided the choice of $k$:

1. **Stability of predictive performance.** We observed that at $k = 40$, the gap between training and test correlations was minimized, indicating reduced risk of overfitting and improved generalizability. Although higher latent dimensions produced comparable raw correlation values, they were accompanied by larger train–test discrepancies, suggesting diminished stability.

2. **Consistency of gene-level influence.** To further assess interpretability, we examined the combined influence (Dependency effect + $IC_{50}$ effect) of the top five and bottom five influential genes across different latent dimensions. At $k = 40$, these genes displayed relatively concordant influence patterns, with less variability compared to other latent settings. This indicates that the model's attribution of gene effects is more stable and biologically interpretable at this dimensionality.

Taken together, these analyses support the selection of $k = 40$ as an optimal compromise: sufficiently large to capture the major sources of variation in the data, yet small enough to ensure stable predictions and interpretable gene-level contributions.

#### FORGE under-performs when using unrelated target in predicting dependency and drug responses

We showed that FORGE performed well in predicting $IC_{50}$ and target gene dependency for numerous drug-target pairs (Ref. Results). We proved the convergence of FORGE through detailed mathematical formulations. Given our results, we now wanted to test our FORGE algorithm if the drug is paired with an unrelated gene. We believe this test would identify whether FORGE learns representations specific to the target gene, and systematically identify trends in benefit scores given a pair of drug and an unrelated gene. To this end, we selected the gene A1CF to be paired with erlotinib. A1CF is known to be expressed in all tissues similar to EGFR, and is one of the key genes in promoting epithelial-mesenchymal transition (EMT), and has been associated with progression of breast and lung cancers [1].

Given that A1CF and EGFR play overlapping biological roles, we first wanted to check whether A1CF's expression is influenced by therapy with erlotinib. For this, we queried the CTR-DB v 2.0 server (`http://ctrdb.cloudna.cn/browse/dataset`) [2]. This database facilitates comparative gene expression among responders and non-responders to a particular therapy by mining multiple studies. For erlotinib, we identified a total of 46 samples tested, among which 43 were from a single micro-array dataset (GEO accession: GSE61676). For the purpose of analysis, we chose arbitrary cutoffs of P and adjusted P values of 1, and absolute $\log_2$-fold change of 0 for getting the full list of genes. We then observed no change in A1CF's expression among the responders when compared with non-responders (fold-change = 0.043, adjusted P = 0.99).

We used the 658 cell lines that were used when analyzing the EGFR-erlotinib pair, and considered the same set of highly-correlated genes (HCGs). While using HCGs specific for the erlotinib-A1CF pair would be the optimal choice, we did not explicitly wanted to generate a model for some unrelated target. Instead, we wanted to check if the original HCGs set is specific to the target pair erlotinib-EGFR, and whether the gene influence scores vary significantly given a new list of dependency values. As expected, we did identify very poor correlation between predicted and actual dependencies and $IC_{50}$ (Pearson $r$ = -0.05 and 0.32 respectively) for the erlotinib-A1CF model (Supplementary figure 9a-b). Similarly, the computed benefit scores failed to capture trends in drug responses and gene essentiality (Supplementary figure 9c-d). We can therefore conclude the model's accuracy is largely determined by the target gene selected for the model, and a random gene may give under-performing model. This single comparison highlights the FORGE's ability to learn for two separate tasks that are biologically linked.

After finding poor performance of FORGE in modeling unrelated drug-target pair, we wanted to assess the gene influence scores that drive the model's performance. We wanted to compare the influence scores for $IC_{50}$ and dependency similar to the approaches described for the erlotinib-EGFR pair. When computed, we observed similar correlation among the influence scores (Spearman $\rho = 0.14$) when compared with the original pair (Spearman $\rho = 0.16$) (Figure 4a). Interestingly, the top 40 genes for the original pair had very low gene influence scores, with most genes having scores close to zero (Supplementary figure 8e).

From the above analyses, we conclude that FORGE performance is maximal when a drug is paired with its own target gene, and the benefit scores by their association with dependency and $IC_{50}$ can differentiate false hits even when underlying biological functions are similar. It should be noted that we tested on a single unrelated drug-target pair, and our findings are limited to the erlotinib-A1CF pair alone. Nevertheless, we believe similar findings can be observed for any other gene when paired with a drug that does not target it.

#### Linearity in Model Building vs. Nonlinearity in Feature Selection

Although the *FORGE* architecture itself is based on a linear joint factorization, which favors interpretability and tractable optimization, the feature selection strategy incorporated a nonlinear element. Specifically, we selected high-confidence genes by evaluating their association with both dependency scores and $IC_{50}$ values using Spearman's rank correlation. Spearman correlation is inherently nonparametric and capable of capturing monotonic but nonlinear relationships between variables.

This design means that while the final *FORGE* model operates linearly in projecting expression data into a shared latent space, the upstream filtering of features already integrates a layer of nonlinearity. In practice, this hybrid approach balances two needs: (i) ensuring that the gene set reflects both linear and nonlinear associations with drug sensitivity, and (ii) maintaining the interpretability and theoretical guarantees of the linear *FORGE* framework.

Thus, nonlinearity is introduced during feature selection, whereas model building itself retains linearity for transparency and stability. This separation reflects a deliberate compromise: nonlinear signals are acknowledged when choosing the input space, but the downstream model remains interpretable and amenable to rigorous validation.

#### Deriving Raw Counts from FPKM: Accurate vs. Approximate

**1. Accurate Formula**
The definition of FPKM for gene $i$ is:

$$\text{FPKM}_i = \frac{\text{Raw Count}_i \times 10^9}{\text{Effective Length}_i \times \text{Mapped Fragments}}. \tag{1}$$

Rearranging gives:

$$\text{Raw Count}_i = \text{FPKM}_i \times \frac{\text{Effective Length}_i \times \text{Mapped Fragments}}{10^9}. \tag{2}$$

Assumptions:

- Average paired-end fragment length $\approx 200$ bp (75–100 bp reads in the original data).
- Effective gene length: $\text{Effective Length}_i = L_i - 199$.
- Total mapped fragments: $35 \times 10^6$.

Substituting these values:

$$\text{Raw Count}_i = \text{FPKM}_i \times \frac{(L_i - 199) \times 35}{1000}. \tag{3}$$

—

**2. Approximate Formula**
For practical computation, we may approximate:

$$\text{Raw Count}_i \approx \frac{\text{FPKM}_i \times L_i}{35}. \tag{4}$$

*Justification.* Starting from the accurate expression:

$$\text{Raw Count}_i = \text{FPKM}_i \times (0.035L_i - 6.965),$$

whereas the approximation yields

$$\text{Raw Count}_i \approx \text{FPKM}_i \times 0.02857L_i.$$

The two expressions are close in magnitude, with the approximation sacrificing a small amount of accuracy for algebraic simplicity.

—

**3. Worked Example**
Consider:

- Gene length: $L_i = 1000$ bp
- FPKM: 10
- Fragment length: 200 bp $\Rightarrow$ Effective length = 801 bp

**Accurate:**

$$\text{Raw Count} = 10 \times \frac{801 \times 35}{1000} = 10 \times 28.035 = \boxed{280.4}$$

**Approximate:**

$$\text{Raw Count} \approx \frac{10 \times 1000}{35} = \boxed{285.7}$$

*Relative error:* $\sim 1.9\%$ overestimation by the approximate formula.

—

**4. Handling Variable Read Lengths (Paired-End)**

When read lengths vary (e.g. 75–100 bp), the fragment length may be approximated as:

$$\text{Fragment Length} \approx 2 \times \text{Read Length} + 50.$$

Thus:

$$\text{Effective Length}_i = L_i - \text{Fragment Length} + 1,$$

which can be substituted into the accurate formula:

$$\text{Raw Count}_i = \text{FPKM}_i \times \frac{\text{Effective Length}_i \times 35}{1000}.$$

—

**5. Summary**

| **Version** | **Formula** | **Remarks** |
|---|---|---|
| Accurate | $\text{FPKM}_i \times \frac{(L_i - 199) \times 35}{1000}$ | Accounts for fragment length and mapped fragments |
| Approximate | $\frac{\text{FPKM}_i \times L_i}{35}$ | Simpler, small bias, ignores effective length |

### References


1. Wang, L. & Cheng, Q. APOBEC-1 Complementation Factor: From RNA Binding to Cancer. *Cancer Control* **31,** 10732748241284952 (2024).
2. Jiang, J. *et al.* CTR-DB 2.0: an updated cancer clinical transcriptome resource, expanding primary drug resistance and newly adding acquired resistance datasets and enhancing the discovery and validation of predictive biomarkers. *Nucleic Acids Research* **53,** D1335–D1347 (2025).


### Supplementary Figures

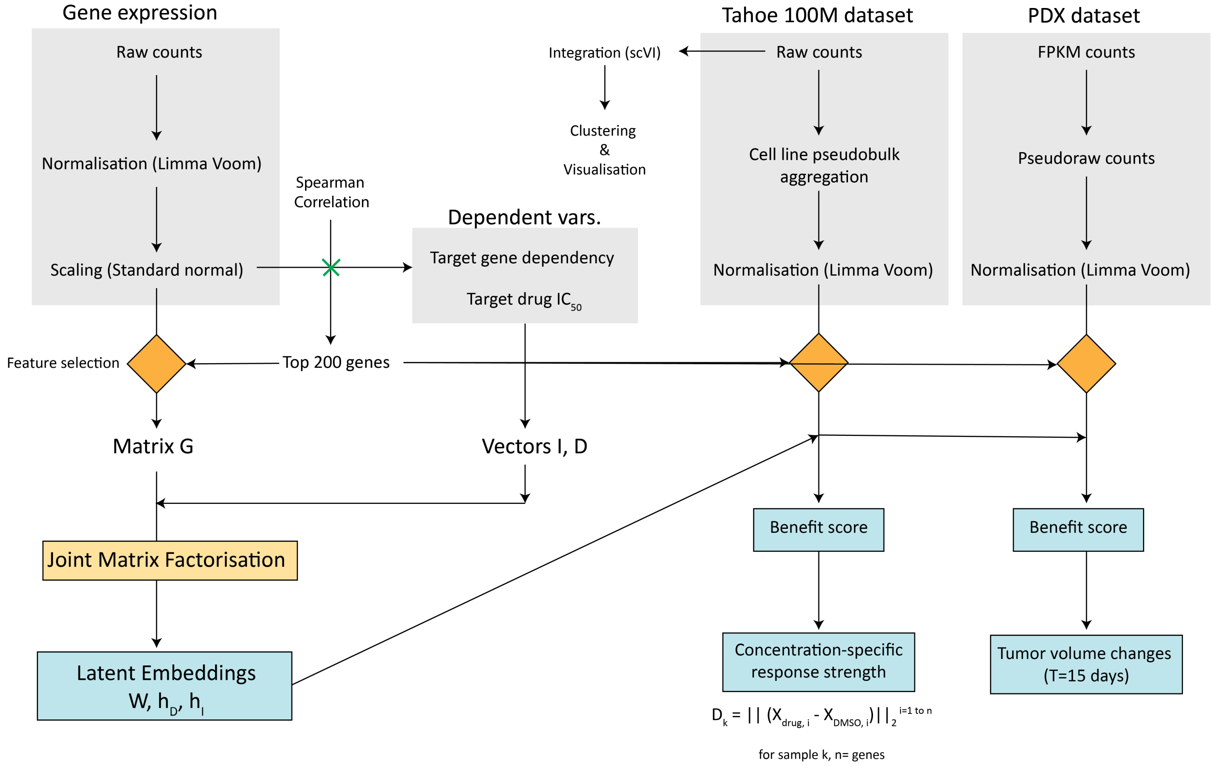


**Supplementary Figure 1. Overview of the preprocessing steps**. Raw counts were normalized using `voom`, after which input features were selected based on their correlation (Spearman's $\rho$) with the target variables—gene dependency and drug $IC_{50}$. The validation datasets included this set of highly correlated genes (HCGs), and Benefit scores were computed for all datasets. Response variables analyzed included percent change in tumor volume for the PDX dataset and response strength for the Tahoe-100M dataset. The associated GitHub repository provides a generalized pipeline for `voom` normalization along with preprocessing steps specific to each dataset.

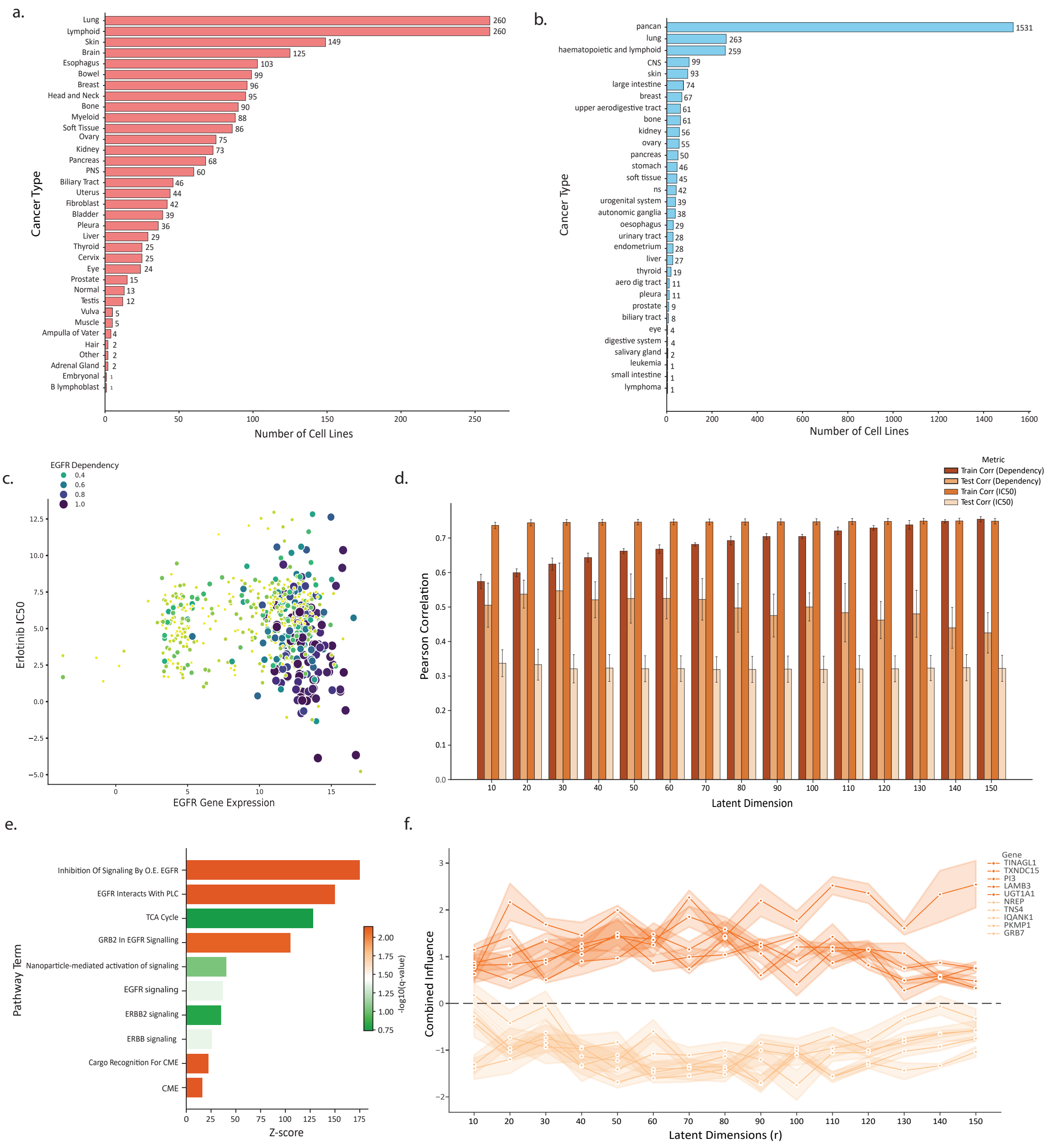


**Supplementary Figure 2. Exploratory analysis of the datasets**. (a-b) Cancer types among the 1034 cancer cell lines in DepMap (a) and 1325 cell lines in CREAMMIST (b) datasets. (c) Bubbleplot depicting EGFR expression levels and erlotinib $IC_{50}$, and bubble sizes represent EGFR dependency. Clearly highly dependent cell lines have high expression and low $IC_{50}$. (d) Tuning the hyper-parameter $k$ for the FORGE model. The error bars represent 95% C.I. for the five bootstrap iterations per instance. (e) Gene set enrichment analysis of top 40 genes with highest/lowest influence scores. EGFR signaling was the top enriched terms (f) Gene importance scores for various instances of FORGE when tuning $k$.

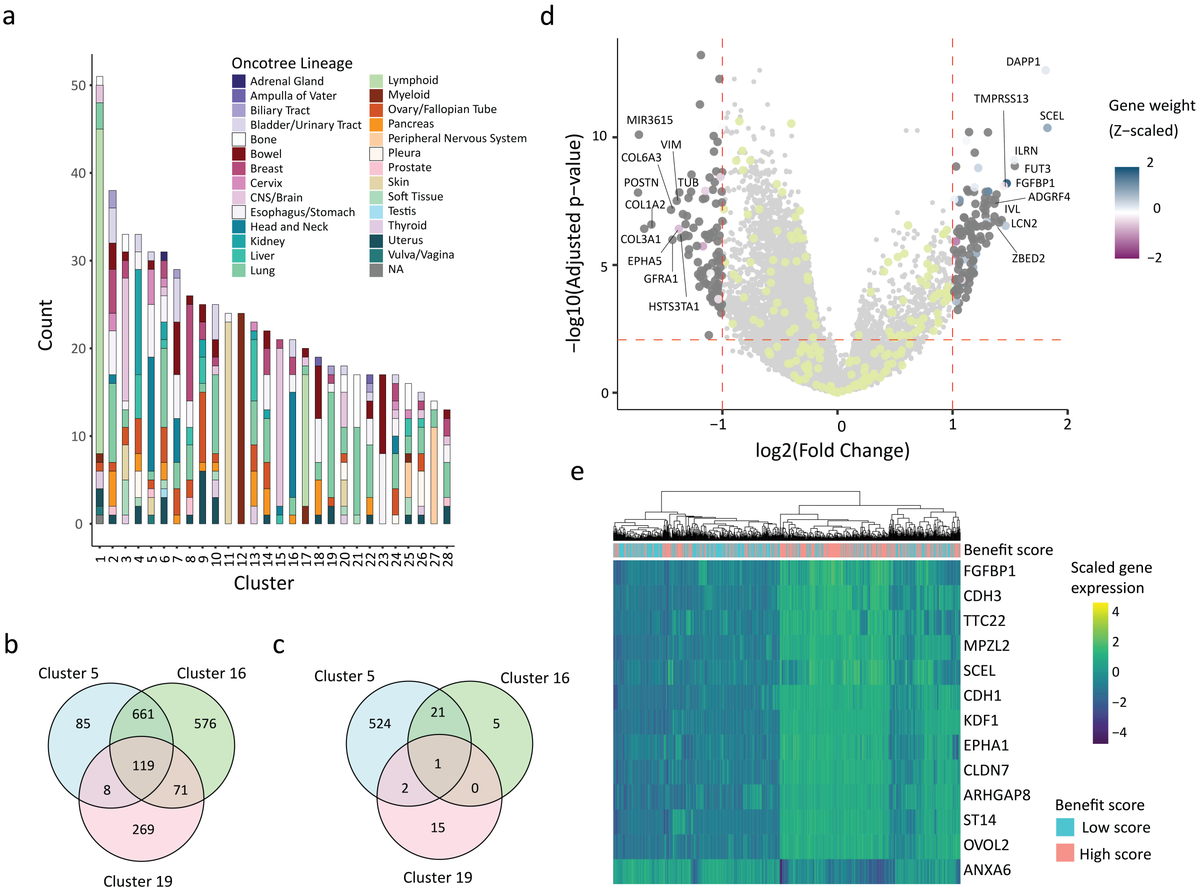


**Supplementary Figure 3. Key findings from highly susceptible clusters**. (a) Stacked barplot showing cluster-wise tissue of origin (Oncotree lineage). One cell line in cluster 1 had no lineage specified (NA). (b-c) Common and unique up (b) and down-regulated (c) genes among the three highly susceptible clusters. (d) Volcano plot depicting the gene expression changes among the 1034 cell lines in DepMap categorized through benefit score groups. The HCGs that are significantly up/down-regulated were colored by their gene weights, while non-significant DEGs are colored in greenish-yellow. (e) Heatmap showing the gene expression levels of 13 genes identified as common DEGs for high benefit scores and highly responding clusters. .

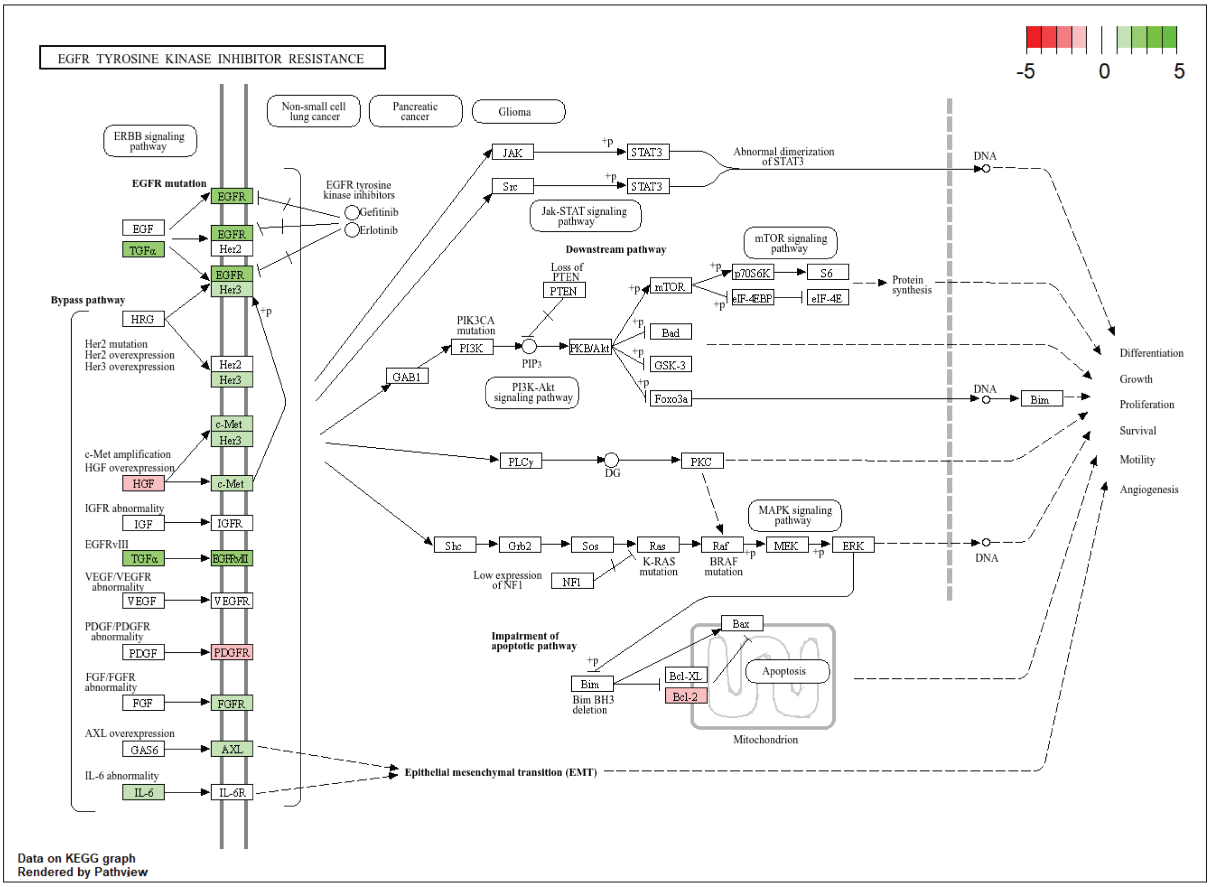


**Supplementary Figure 4. KEGG Pathway changes associated with gene signature for susceptible clusters**. KEGG pathway map enrichment for EGFR-Tyrosine kinase inhibitor resistance. The upregulated genes are in green and repressed genes are in red. The image was generated using PathView in R for the KEGG pathway HSA:01521 (EGFR tyrosine kinase inhibitor resistance)

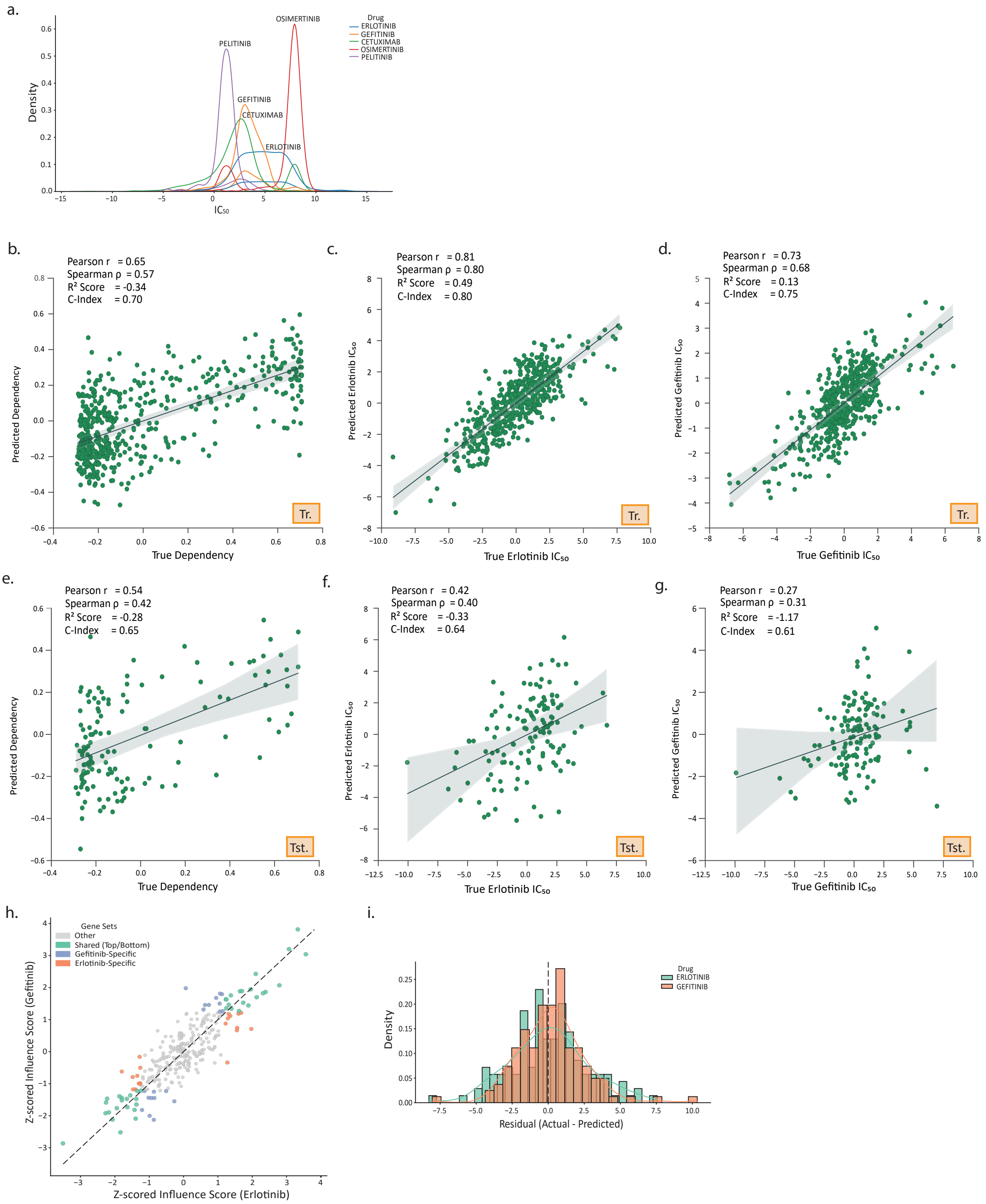


**Supplementary Figure 5. Performance of the Multi-drug FORGE implementation.** (a) Density plot showing the $IC_{50}$ values of multiple drugs targeting EGFR. Only erlotinib and gefitinib have high variance favouring modelling approaches. (b-d) and (e-g) train and test performances of the multi-drug FORGE in predicting dependency (b, e), erlotinib $IC_{50}$ (c,f) and gefitinib $IC_{50}$ (d,g). (h) Observed gene influence scores for erlotinib and gefitinib in the multi-drug model. (i) Histogram showing the distribution of $IC_{50}$ residuals by the model. The residuals are near-normal indicating absence of overfitting.

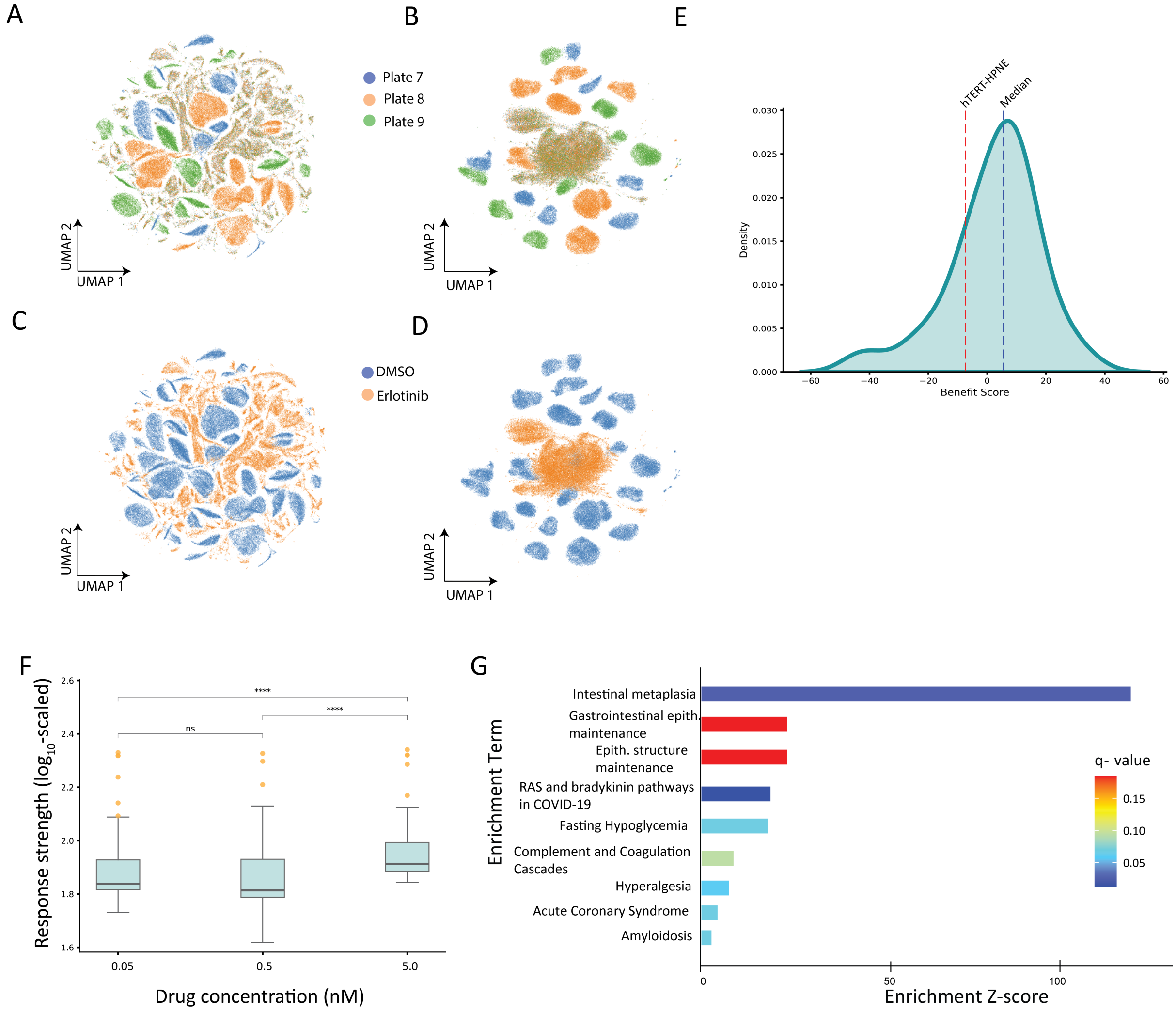


**Supplementary Figure 6. Analysis of the Tahoe-100M dataset.** (A-D) UMAP plots before and after integrating with scVI: before (A) and after (B) integration of multiple batches colored by batch IDs (plates), and before (C) and after (D) integration of multiple batches colored by treatment categories - DMSO or erlotinib. Only the highly responding cell-lines were shown. E) Density plot depicting the distribution of Benefit Scores computed for all the cell lines. The median value and the Benefit score for normal (hTERT-HPNE) are shown for reference. F) Concentration-specific response strengths for all cell lines. Significantly higher response strengths were noted at high (5.0 nM) concentrations. G) Gene-set enrichment analysis using enrichr-KG with relaxed filtering criteria - gene maps to at least one term, each term has at least three genes, q-value threshold set at 0.1. *Significance codes:* **** $p < 0.0001; n.s., not significant.$

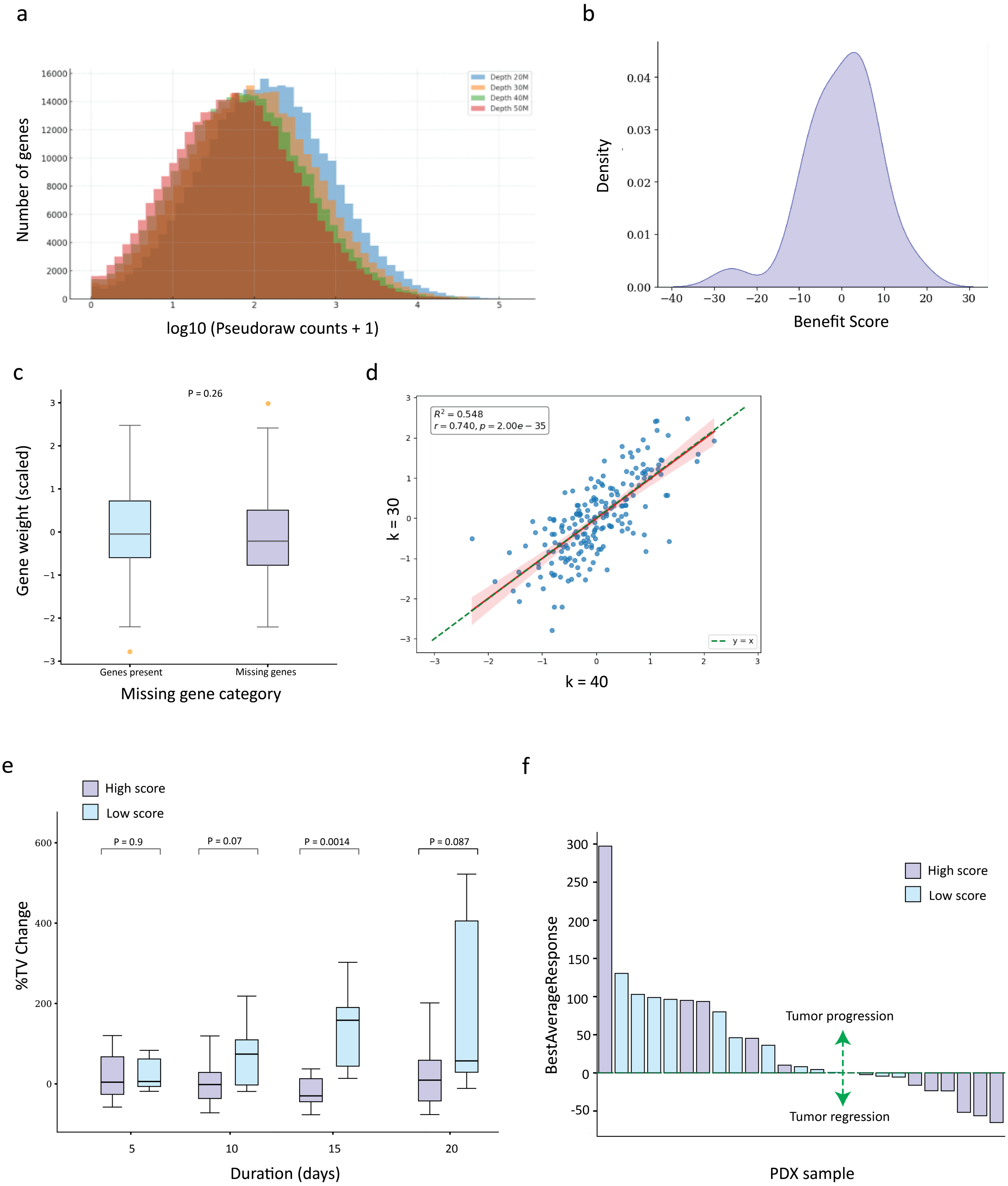


**Supplementary Figure 7. Analysis of the PDX dataset.** (a) Distribution of the pseudo-raw counts for 50,000 genes with randomly initiated FPKM values and different depths (20-50M mapped fragments). Overall, the distributions are similar with shift in the mean counts. (b) Density of the Benefit Scores for the 25 samples included in the analysis. Note that the shape of the curve resembles that of Tahoe-100M (Supplementary figure 6E). (c) Weights for the genes present and absent in the PDX dataset. The weights were not significantly different (P=0.26), revealing a loss of information due to these missing genes. (d) Scatterplot depicting weights obtained through k=30 and k=40 settings in FORGE model. Overall the weights were similar and had high correlation (Pearson's $r = 0.74$) (e) Boxplot depicting percent change in tumor volumes (%TVChange) binned in durations of five days. Significant differences were observed at T=15 days. Note that very few samples had measurements beyond 15 days. (f) Waterfall plot depicting the Best average response for all the 25 patients. If the tumor is repressed due to therapy - the %TVChange and best-average-responses are negative.

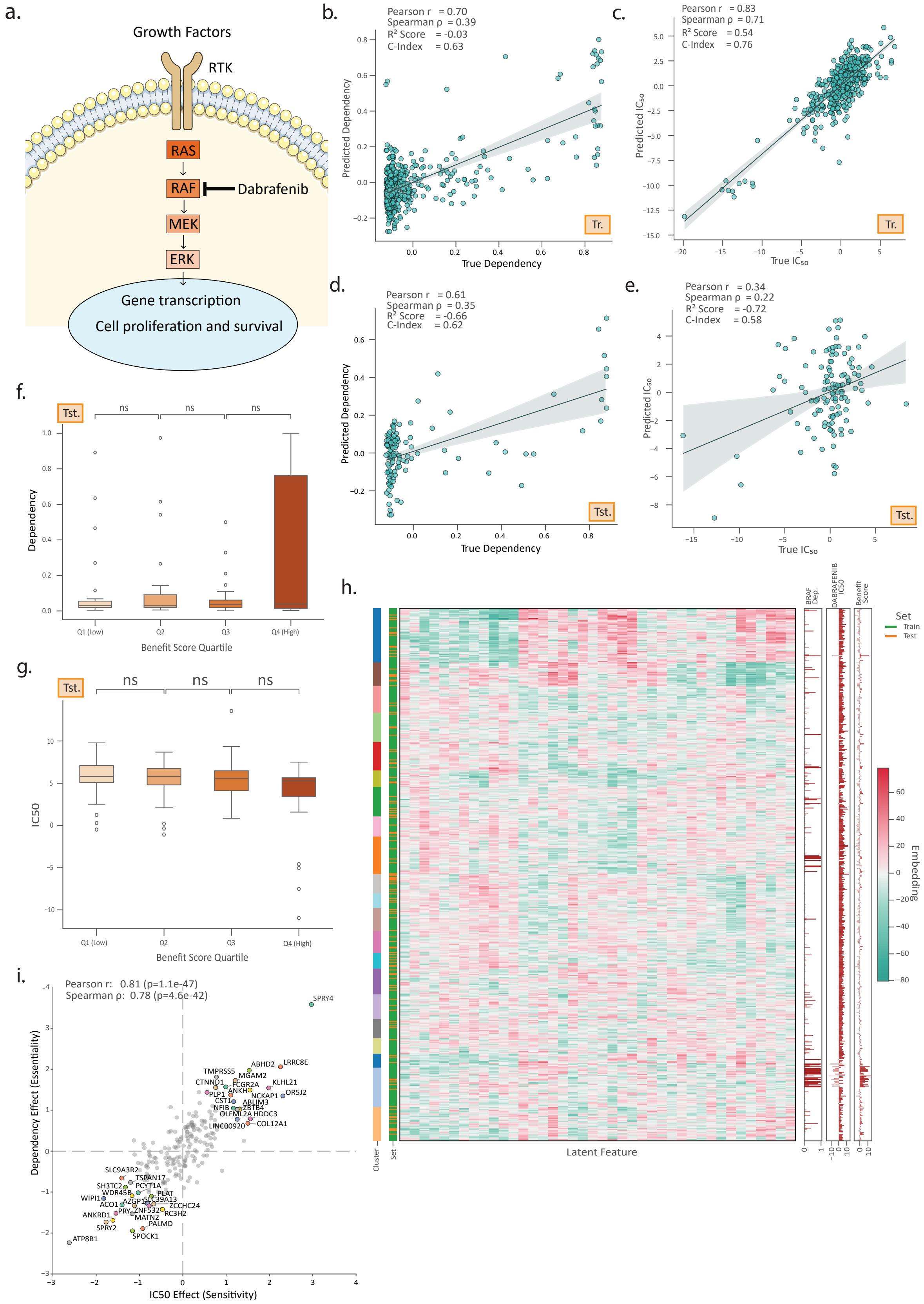

**Supplementary Figure 8. Performance of the FORGE model for the Dabrafenib–BRAF drug–target pair.** (a) Schematic of Dabrafenib mechanism of action in the MAPK signaling cascade. (b-c) Scatterplot of true vs. predicted BRAF dependency values. (d-e) Scatterplot of true vs. predicted Dabrafenib $IC_{50}$ values. (f-g) Quartile boxplots of dependency and $IC_{50}$ values across Benefit Score quartiles in test data. (h) Heatmap of latent embeddings showing 21 distinct clusters. (i) Gene influence scatterplots showing top and bottom influencer genes for dependency and $IC_{50}$ effects.

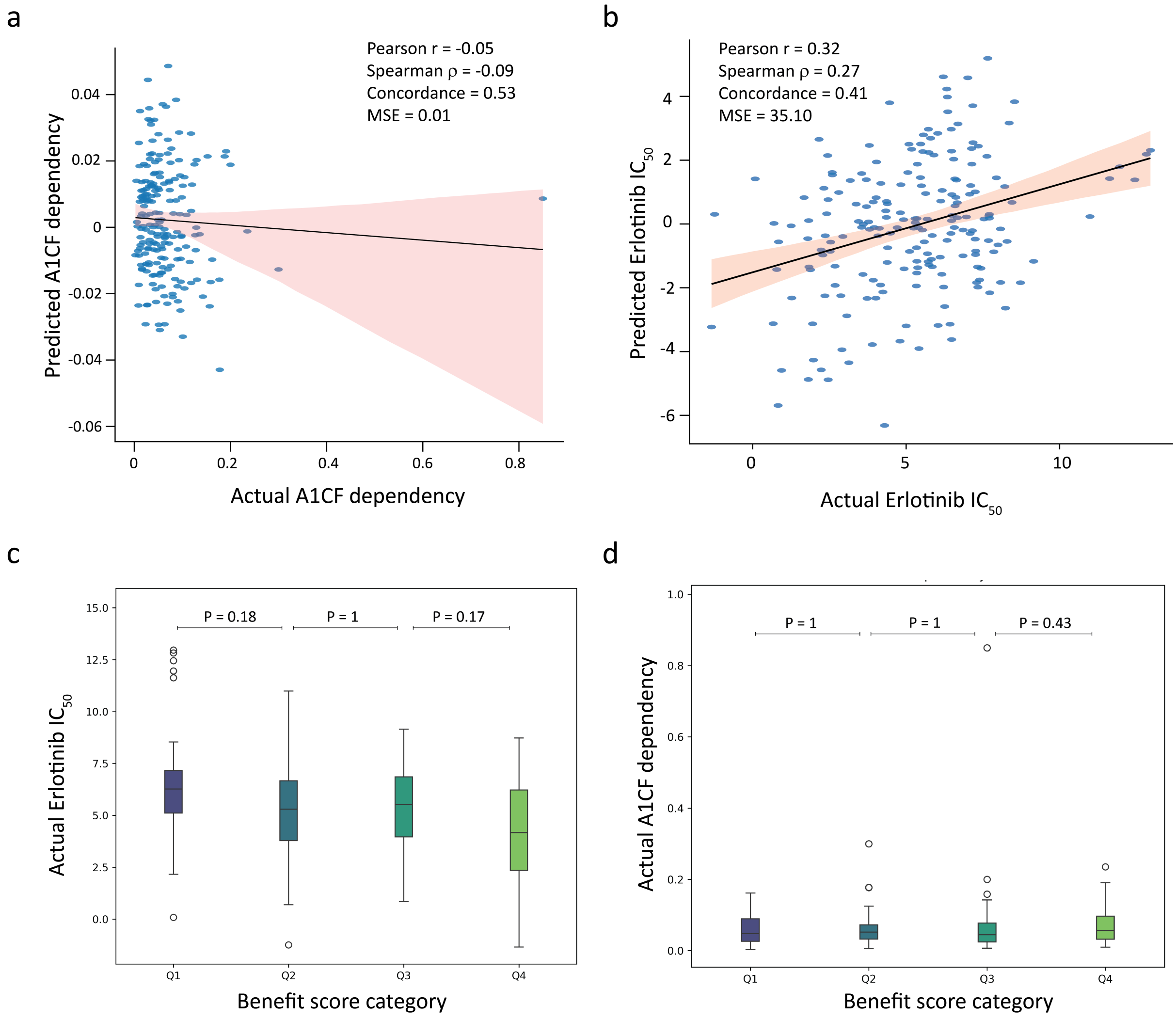


**Supplementary Figure 9. FORGE when tested on unrelated target gene A1CF.** (a-b) Scatterplot denoting correlation and concordance of actual and predicted values for dependency (a) and $IC_{50}$ (b). The plots are for the test dataset. (c-d) Boxplots showing quartile-wise segregation of benefit scores (X-axis) and $IC_{50}$ and gene dependency. P values were computed using two-sided Mann-Whitney test (e) Scatterplot showing gene influence scores with key genes from the EGFR-erlotinib pair marked. The values have poor correlation when compared with the related target (see Figure 4a)

### Supplementary tables

**Table 1:** Count of Primary Or Metastasis among key clusters

| Cancer type | Key clusters | Other clusters | Total |
|---|---|---|---|
| Primary | 43 | 331 | 374 |
| Metastatic | 23 | 214 | 237 |
| **Total** | **66** | **546** | **612** |

46 cell lines had no recorded metadata for cancer types.
